# Nuclear morphology and chromatin organization modulate T cell cytoskeletal remodeling and immune synapse formation

**DOI:** 10.1101/2025.08.25.672234

**Authors:** Ivan Rey-Suarez, Aashli Pathni, Frank Fazekas, Matthew Connell, Arpita Upadhyaya

## Abstract

T cell activation is characterized by rapid reorganization of the actin cytoskeleton and cell spreading on the antigen presenting cell. The T cell nucleus occupies a large fraction of the cell volume, and its mechanical properties are likely to act as a key determinant of activation. However, the contribution of nuclear mechanics to T cell spreading and activation is not well understood. Mechanical rigidity of lymphocyte nuclei is conferred by chromatin compaction and dense packing of heterochromatin. We find that nuclear deformation and increased chromatin compaction accompany T cell spreading, in T cells. Reducing chromatin compaction leads to increased cell spread area and nuclear deformation, while diminishing accumulation and peripheral enrichment of F-actin at the immune synapse. In contrast, enhanced chromatin compaction reduced spread area and nuclear deformation, which was accompanied by increased peripheral F-actin organization at the immune synapse. These findings suggest a reciprocal interaction between chromatin compaction and actin cytoskeletal organization. We identified SUN proteins and myosin as critical elements through which chromatin compaction orchestrates actin morphology and cell shape, facilitating T cell adaptation to antigen-presenting surfaces of varying stiffness. These results emphasize the crucial role of chromatin compaction in T cell activation, underlining the mechanical relationship between the nucleus and the cytoskeleton during immune responses, and suggest new avenues for understanding T cell mechano-responsiveness.

## Introduction

Upon recognition of antigen at the surface of an antigen presenting cell (APC), T cells spread over the APC, increasing their contact area and enhancing cell-cell communication^1, 2^. During this process of T cell activation, the actin cytoskeleton is dramatically remodeled, and the microtubule cytoskeleton becomes polarized, with the centrosome translocating towards the cell-cell contact^3–5^. Cell spreading is accompanied by extensive reorganization of signaling molecules to ultimately form a distinct structure called the immune synapse, which is essential to maintain sustained signaling required for T cell activation^6, 7^.

The nucleus occupies a significant portion of the cytoplasmic space in T cells^8^. As is well known for adherent cells on artificial substrates, spreading due to actin polymerization results in the movement of the nucleus towards the plane of the substrate as well as a flattening of the nucleus^9, 10^. Given the high rigidity of the nucleus^11^, changes in cell shape are constrained by changes in nuclear shape and consequently by the mechanical properties of the nucleus. Nuclear deformations result in the reorganization of chromatin and import of signaling molecules and transcription factors, thus serving as a platform for gene expression that alters cell states and promotes cellular adaptation to environmental surroundings^12^. The impact of nuclear mechanical properties on the evolution of cellular morphology of T cells during activation is not well understood.

The mechanical behavior of the structural elements in the nucleus is crucial to understanding nuclear deformation in cells. The two main components of mammalian nuclei that determine their mechanical properties are the nuclear lamina and the chromatin^13^. However, T cells contain limited lamin A/C prior to activation, and lamin A/C levels increase over days following activation^14, 15^. Chromatin, which fills the nucleus, is a repository of genetic information and imparts structure and mechanical rigidity to the nucleus. The epigenetic state and the three-dimensional organization of chromatin and distribution of the nucleoplasm govern the elastic properties of the nucleus^16–18^.

Chromatin in resting or naive T cells is enriched in heterochromatin, which is densely packed and consists of transcriptionally repressed genes and methylated histones in close proximity to the nuclear envelope. Heterochromatin is relatively stiff and compacted, and the loss of heterochromatin has been shown to soften the nucleus^16, 17, 19^. Lymphocyte activation results in extensive chromatin remodeling both at early (within minutes to hours) and over longer timescales (days), leading to transcriptomic changes that tune the cellular state^8, 19^. These changes also alter the mechanical properties of the nucleus and consequently, its morphology. The nucleus is physically coupled to the cytoskeleton via the LINC (Linker of Nucleoskeleton and Cytoskeleton) complex, which consists of SUN and KASH domain proteins that span the nuclear envelope and create a direct mechanical connection between nuclear and cytoplasmic compartments, relaying forces generated by cytoskeletal changes to the nucleus^20^. Thus, the degree of chromatin compaction and its corresponding contribution to nuclear mechanical properties is likely to modulate cytoskeleton morphology and the spreading of cells during the initial phase of T cell activation, with implications for subsequent stages of the T cell response^21^.

Given the importance of rapid, cytoskeletal driven spreading of T cells to form the immune synapse^2, 6^, we examined how nuclear morphological changes influence T cell spreading and early activation. We used pharmacological treatments and RNA interference to tune chromatin compaction and characterized the resulting changes induced in cytoskeletal morphology and dynamics. We then examined the influence of chromatin compaction on T cell mechano-responsiveness. We identified SUN proteins and myosin motors as key elements through which chromatin compaction modulates actin morphology and cell shape and allows T cells to respond to antigen presenting surfaces of different stiffness. Our studies highlight how T cells adjust their nuclear morphology to compressive forces generated by the actomyosin cortex and in turn, how the mechanical properties of the nucleus drive cytoskeletal rearrangements at the immune synapse and thus enable T cells to adapt to their mechanical surroundings and maintain T cell activation.

## Results

### T cell signaling activation induces nuclear deformation and changes in chromatin compaction

We first quantified the morphological dynamics of the T cell nucleus during activation. Jurkat T cells transiently transfected with tdTomato-F-tractin (to label F-actin) and stained with SiR-DNA (to label the nucleus) were allowed to spread on anti-CD3 coated coverslips. Spinning disk confocal microscopy was used to visualize cells in 3D during activation (Figure 1A). We observed that the actin cytoskeleton reorganized to form a characteristic ring at the synapse (Figure 1A upper panels) during the first few minutes of spreading^22^. As the cell spread, the cell and nucleus height decreased over time (Figure 1A lower panels), leading to a change in the z-position of the nucleus centroid (Figure 1B). To parametrize the change in nucleus shape over time, we defined the nuclear deformation ratio as the ratio between the height and average x-y radius of the nucleus. During cell spreading, the nucleus is compressed and expands parallel to the glass surface and reaches maximum deformation at ∼ 5 minutes after initiation of activation (Figure 1C). To characterize changes in nuclear morphology, we computed the nucleus surface (Figure 1D) using the segmented chromatin signal from confocal stacks (Supplementary Figure 1A). We found that the change in nuclear shape is accompanied by a small transient change in volume while maintaining a constant surface area (Supplementary Figure 1B-C).

**Figure 1.**
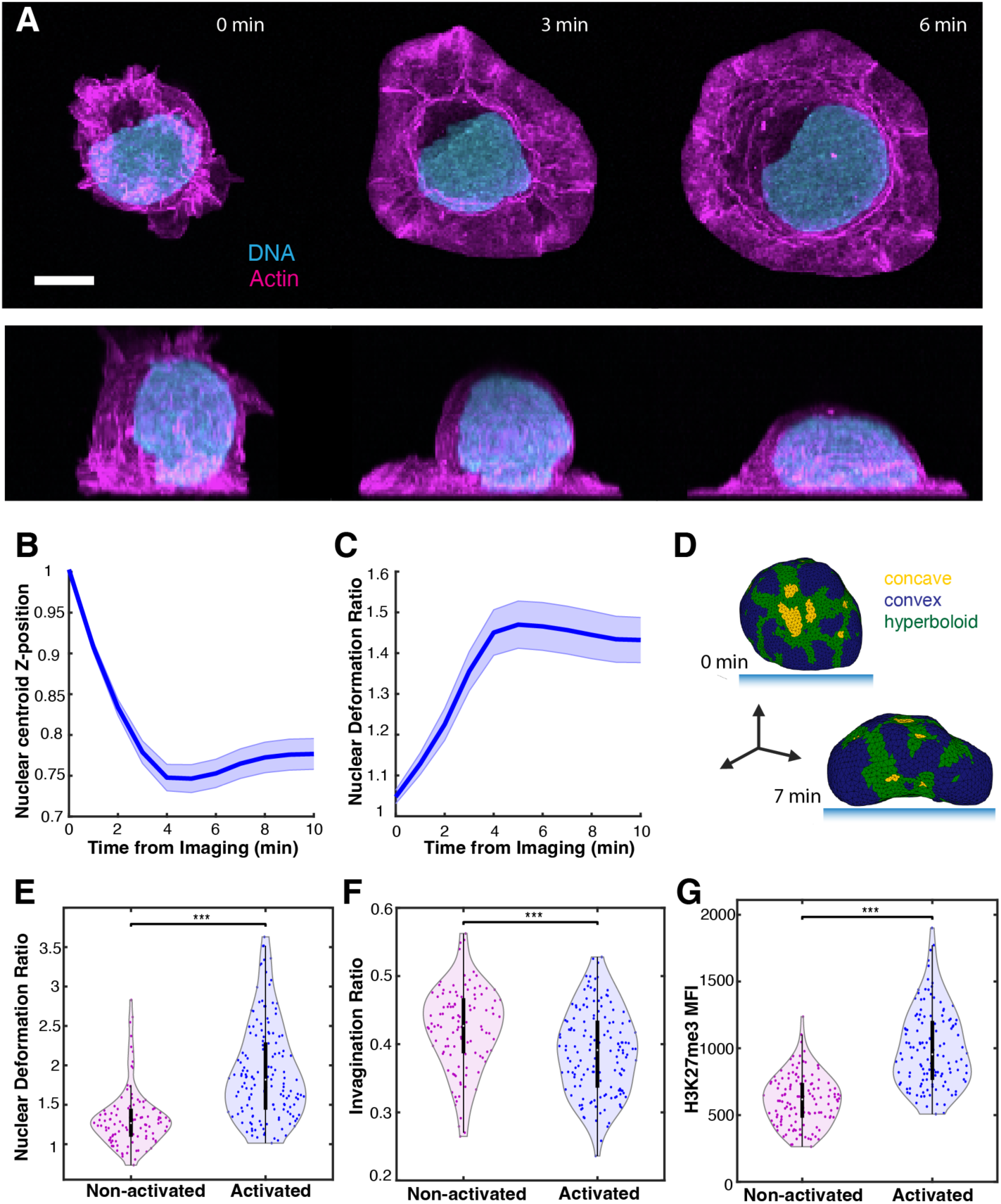
Activation induces significant morphological changes in the T cell nucleus. A) Maximum intensity projections in *xy* (upper panels) and *xz* (lower panels) of a tdTomato-F-tractin (magenta) expressing Jurkat T cell stained with SiR-DNA (cyan) as it spreads on a coverslip coated with anti-CD3 at 0 (left), 3 (center) and 6-minute timepoints (right). B) Normalized change in the nucleus centroid Z-position as a function of time. Shaded portion of the curve is the standard error (*n*=69 cells). C) Plot of the nuclear deformation ratio (ratio of height to average max radius) as a function of time. Shaded portion of the curve is the standard error (*n*=69 cells). D) Typical mesh model obtained from the segmented 3D images of the cell nucleus; green sections correspond to hyperboloid regions and yellow sections to concave regions of the nuclear surface. E) Comparison of the deformation ratio for activated cells versus non-activated cells (*p*<0.001 Wilcoxon rank sum test, *n* = 109 and 150 cells respectively). F) Plot of the invagination ratio for activated versus non-activated cells (*p*<0.001 Wilcoxon rank sum test, *n* = 109 and 150 cells respectively). G) Plots of mean fluorescence intensity (MFI) for the histone methylation mark, H3K27me3 (*p*<0.001 Wilcoxon rank sum test, *n*=130 cells for both conditions). Scale bar corresponds to 5 μm.

To study differences in nuclear morphology induced by activation, we fixed cells after seven minutes of being seeded onto anti-CD3 coated (activated) or poly-L-Lysine (PLL) coated (non-activated) coverslips. Activated cells displayed significantly higher nuclear deformation ratios than non-activated cells (Figure 1E). The surface of the nucleus exhibited a complex topography.

To quantify the surface, we calculated regions with hyperboloid, concave and convex curvature (colored green, yellow and blue, respectively; Figure 1D, Supplementary Figure 1A-B). We defined invaginations as regions with negative curvature and the invagination ratio as the ratio of the area of concave plus hyperboloid regions to the total nucleus surface area (see Methods). Activated cells displayed significantly lower invagination ratios than non-activated cells (Figure 1F), indicating that activation smoothens the nuclear surface. To test whether primary T cells also exhibited nuclear deformations, we activated murine CD8+ T lymphocytes (cytotoxic T lymphocytes or CTLs) on anti-CD3 coated coverslips. Upon activation, the nucleus moved towards the substrate and deformed. The time course and relative extent of nuclear deformation was similar to that observed in Jurkat T cells (Supplementary Figure 1D-F). Furthermore, this deformation was dependent on activation. as activated CTLs showed increased nuclear deformation compared to non-activated cells (Supplementary Figure 1G).

Transient nuclear deformation has been shown to induce global changes in chromatin compaction in fibroblasts^23^. We investigated whether nuclear shape change during T cell spreading led to changes in chromatin compaction. We used immunostaining to determine the levels of tri-methylation of histone 3 on lysine residue 27 (H3K27me3), a known marker of facultative heterochromatin (compact chromatin), in activated and non-activated Jurkat T cells. We observed that heterochromatin was concentrated mainly at the periphery of the nucleus (Supplementary Figure 2A) and that activated cells displayed a significant increase in H3K27me3 levels compared to non-activated cells (Figure 1G, Supplementary Figure 2B), suggesting an increase in chromatin compaction. We note that the increase in chromatin compaction upon activation is likely to be transient. Previous studies found that T cell activation induced an overall chromatin decondensation via histone acetylation and demethylation^24–26^ after 24-48 hours, facilitating cell proliferation.

### Chromatin decompaction through EZH2 inhibition induces changes in nuclear and actin morphology

The extent of nuclear deformations observed during T cell activation likely depends on its mechanical properties. The major structural elements that determine the mechanical properties of the nucleus are the nuclear lamina and the interior chromatin^13, 17^. Given our observations of changes in heterochromatin distribution and levels, and the known role of chromatin in establishing nuclear rigidity^18, 19, 27^, we hypothesized that the histone modification state of chromatin may contribute to the observed nuclear deformations during activation.

We used DZNep, which inhibits the histone methyltransferase, Ezh2^28, 29^ to reversibly and globally reduce histone methylation, thereby inducing chromatin decompaction. We incubated T cells in DZNep, fixed them after seven minutes of activation on anti-CD3 coated coverslips and immunostained for H3K27me. We found that cells treated with DZNep had lower levels of H3K27me3 than vehicle control cells (Figure 2A, B) as expected. The lower chromatin compaction of DZNep-treated cells did not affect the phosphorylation of Zap70 (Supplementary Figure 3A), an early activation signal, nor did it influence centrosome repositioning towards the synapse (Supplementary Figure 3B), a hallmark of successful T cell activation. These indicate that DZNep treatment preserves T cell signaling and activation over our timescales of interest. Given the role of chromatin modifications in nuclear mechanical compliance, DZNep treated cells should have a lower nuclear stiffness as observed in previous studies^13, 30^. Accordingly, we found that DZNep increased the nuclear deformation ratio (Figure 2C, D) and the invagination ratio (Figure 2E) of activated cells, consistent with nuclear softening.

**Figure 2.**
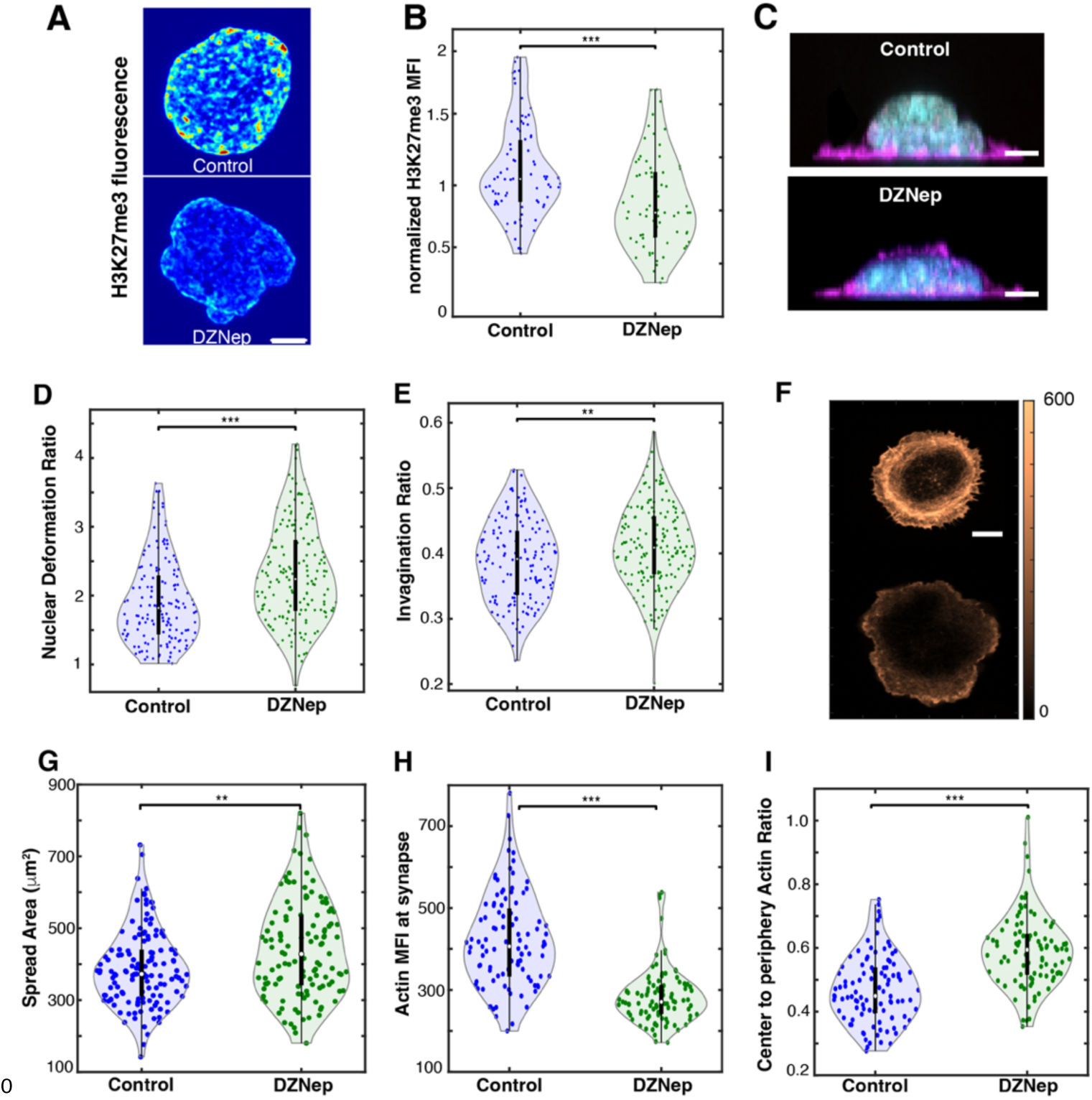
Chromatin decompaction leads to enhanced nuclear deformation and alters synaptic actin morphology. A) Representative fluorescence images of the histone methylation mark H3K27me3 in activated Jurkat T cells with vehicle or DZNep treatment. Both panels are on the same color scale. B) Quantification of H3K27me3 fluorescence intensities for vehicle treated (blue) or DZNep treated cells (green) (*p*<0.001 Wilcoxon rank sum test, *n*=83 and 73 cells respectively). C) Maximum intensity projections in *xz* of activated control and DZNep-treated cells labeled with Hoechst to visualize the nucleus (cyan) and phalloidin to visualize actin (magenta). D) Plots of nuclear deformation ratio for cells treated with DZNep (green) and control (blue) (*p*<0.001 Wilcoxon rank sum test, *n*=150 and 170 cells respectively). E) Invagination ratios for control versus DZNep treated cells (*p*<0.01 Wilcoxon rank sum test, *n*=150 and 170 cells). F) Representative confocal image (synapse plane) showing actin morphology in control and DZNep treated cells stained with phalloidin. Both images are shown with the same color scale. G) Plots of the spread area of control and DZNep treated cells. H) Plots showing the difference in actin accumulation i.e. mean fluorescence intensity (MFI) at the synapse plane for control and DZNep cells (*p*<0.01 Wilcoxon rank sum test, *n*=100 and 99 cells respectively). I) Plots showing the difference in actin organization for control and DZNep treated cells (*p*<0.01 Wilcoxon rank sum test, *n*=100 and 99 cells respectively). Scale bars correspond to 5 μm.

Cell spreading is believed to be sufficient to drive nuclear deformation under a range of conditions^9, 10, 31^ Additionally, the reduction of chromatin compaction and nuclear stiffness has been observed to induce an increase in actin polymerization in human breast cancer cells^32^. Given the connection between nuclear shape deformation, actin polymerization and cell spreading, we next sought to determine whether actin morphology in T cells is altered due to changes in chromatin compaction. We labeled F-actin in fixed cells with Acti-stain 670 phalloidin and used confocal microscopy to investigate the effect of DZNep treatment on cytoskeletal morphology (Figure 2F). DZNep treatment affected synapse morphology, with DZNep-treated cells displaying larger contact areas (Figure 2G), lower actin accumulation at the synapse (Figure 2H) and a smaller degree of actin enrichment at the cell periphery i.e. less pronounced actin rings (Figure 2I).

### HMGA1 knockdown increases chromatin compaction and modulates nuclear deformation and cytoskeleton morphology

Given the effects of decrease in chromatin compaction on T cell morphology, we next investigated the effect of increasing chromatin compaction and correspondingly nuclear stiffness. HMGA1 is a high mobility group protein with high affinity for AT rich regions of DNA, the same regions that are bound by histone H1^33–35^. Loss of HMGA1 has been shown to promote chromatin compaction and increase nuclear stiffness by altering H1 histone distribution and expression^34^. To increase chromatin compaction, we used siRNA-mediated knockdown of HMGA1, validated using immunofluorescence (Supplementary Figure 4A, B). Transfected cells were activated on anti-CD3 coated coverslips, fixed and stained after 7 minutes of activation and imaged with confocal microscopy (Figure 3A). HMGA1 knockdown (siHMGA1) cells, displayed increased H3K27me3 signal (Figure 3B) indicating an increase in chromatin compaction compared to cells transfected with control siRNA. We verified that the change in chromatin compaction did not affect Zap70 phosphorylation (Supplementary Figure 4C) or centrosome repositioning (Supplementary Figure 4D), confirming that T cell activation is unaffected by HMGA1 knockdown.

**Figure 3.**
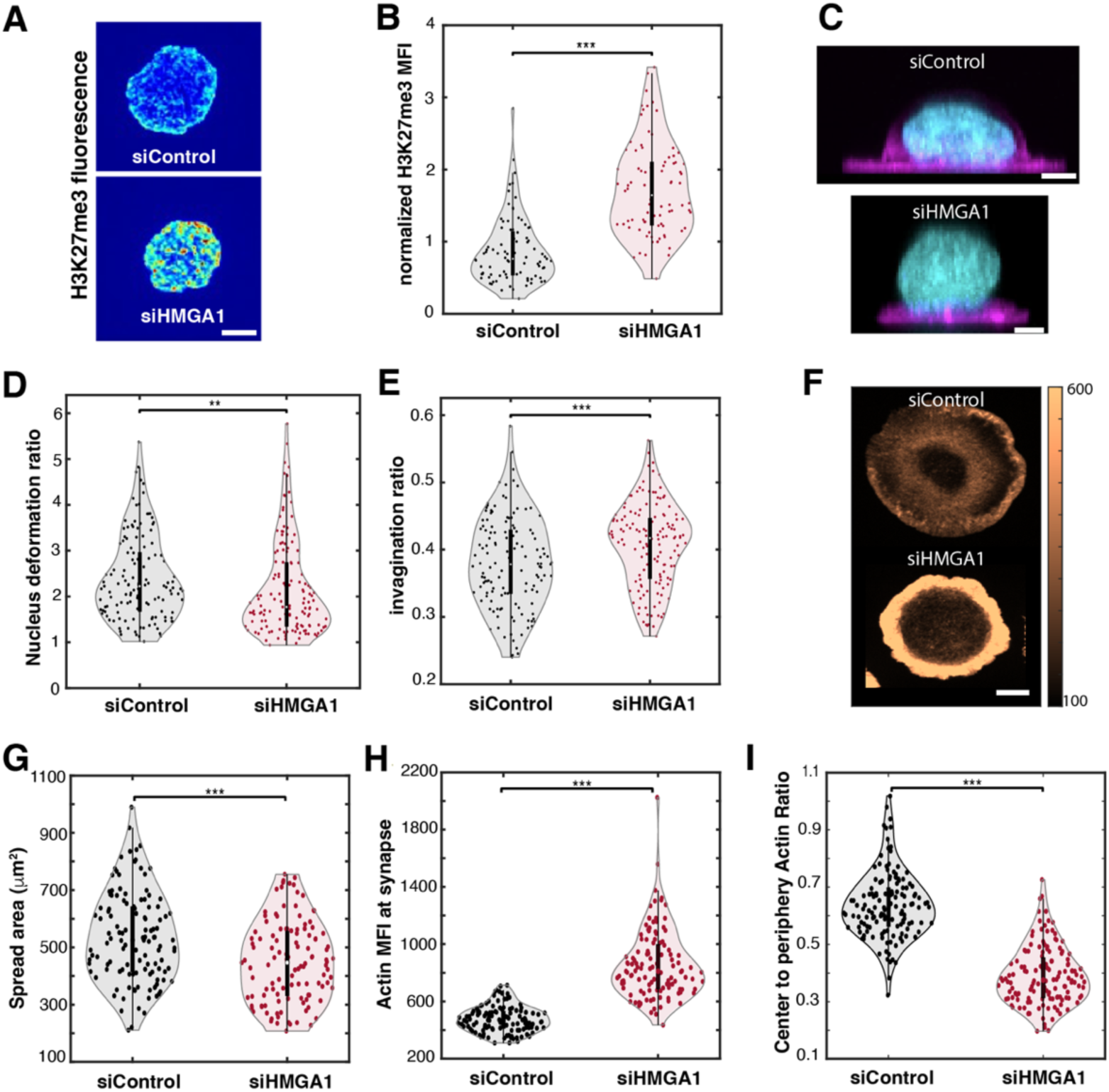
Increased chromatin compaction leads to reduced nuclear deformation and enhancement of synaptic actin. A) Representative images of mean fluorescence intensity of the methylation mark H3K27me3 in activated Jurkat T cells transfected with control silencing RNA (siControl) and cells transfected with HMGA1 silencing RNA (siHMGA1). Both panels are shown on the same color scale. B) Plots showing the mean fluorescence intensity of the methylation mark H3K27me3 in control and HMGA1 knockdown cells (*p*<0.001 Wilcoxon rank sum test, *n*=83 and 84 cells respectively). C) Maximum intensity projections in *xz* of activated cells in siControl or siHMGA1 cells labeled with Hoechst to visualize the nucleus (cyan) and phalloidin to visualize actin (magenta). D) Plots of nuclear deformation ratio for siControl and siHMGA1 cells (*p*<0.01 Wilcoxon rank sum test, *n*=135 and 141 cells respectively). E) Invagination ratios for siControl and siHMGA1 cells. F) Representative confocal image (synapse slice) showing the morphology of actin in control and HMGA1 KD cells labeled with phalloidin. Both images are shown with the same color scale. G) Plots showing the difference in spread area for siControl and siHMGA1 cells (*p*<0.001 Wilcoxon rank sum test, *n*=122 and 117 cells respectively). H) Plots showing the difference in actin accumulation - mean Fluorescence Intensity (MFI) at the synapse slice for siControl and siHMGA1 cells (*p*<0.001 Wilcoxon rank sum test, *n*=135 and 140 cells respectively). I) Plots showing the difference in actin distribution for siControl and siHMGA1 cells (*p*<0.001 Wilcoxon rank sum test, *n*=135 and 140 cells respectively). Scale bars are 5 μm.

Increase in chromatin compaction has been reported to induce an increase in nuclear stiffness^18^. Accordingly, we found that the nuclei of activated siHMGA1 cells had lower deformation ratios than siControl cells (Figure 3C, 3D). Interestingly, we found that the invagination ratios of siHMGA1 cells were also higher than control cells (Figure 3E). This may potentially be explained by the fact that since there is less nuclear deformation in siHMGA1 cells, excess surface area of the nucleus may be partially retained within invaginations. We next examined the effect of increased chromatin compaction on actin morphology in activated cells fixed at 7 minutes post activation and labeled with phalloidin (Figure 3F). We found that siHMGA1 cells displayed lower spread area (Figure 3G), higher actin accumulation at the cell-substrate contact (Figure 3H) and increased actin recruitment to the cell periphery (Figure 3I). Interestingly, the increase in actin accumulation was only observed in activated cells (Supplementary Figure 4A). These observations show that chromatin compaction and concomitant nuclear stiffening leads to reduced cell spreading, but overall higher actin levels and enhanced actin rings at the immune synapse. Taken together, these findings suggest that changes in mechanical properties of the nucleus resulting from chromatin modifications alter cytoskeletal organization at the immune synapse.

### Chromatin compaction state alters actin dynamics

T cell spreading is accompanied by the formation of a peripheral actin ring enriched in Arp2/3 mediated branched actin meshworks, formin-mediated actomyosin arcs adjacent to it and a central region with sparse actin ^36–38^. To determine whether the effects of chromatin compaction on actin morphology and accumulation at the synapse also lead to changes in actin dynamics, we performed TIRF imaging of live tdTomato-F-tractin expressing T cells (DZNep-treated or siHMGA1 and respective controls) activated on anti-CD3 coated coverslips. Timelapse movies were acquired and actin dynamics were analyzed over a 5 minute time window after the cell was maximally spread. To quantify actin dynamics, a square region of interest (ROI) of 2.5 μm was selected at the center and at the periphery (Figure 4A). The fluorescence signal in the ROI was obtained for each timepoint, and the temporal autocorrelation was computed by comparing the intensity value of the first frame with itself and consecutive frames (Figure 4B) and fitting to a double-exponential function^36^ to obtain a measure of the timescale of actin dynamics (see Methods). The correlation decay time was similar for central and peripheral regions (Figure 4C) in siControl cells. In contrast, correlation decay times were significantly larger at the periphery than at the center (Figure 4D) for siHMGA1 cells, suggesting a more stable actin network at the periphery upon increasing chromatin compaction. This is consistent with the increase in actin intensity and enhanced peripheral ring formation in siHMGA1 cells, consistent with more enriched and stable actin morphology. However, DZNep-treated cells displayed similar correlation times at the center and periphery, indicating similar actin dynamics throughout the synapse (Figure 4E). These observations highlight the fact that chromatin compaction can modulate the spatial distribution, stability and dynamics of the actin network in different regions of the immune synapse.

**Figure 4.**
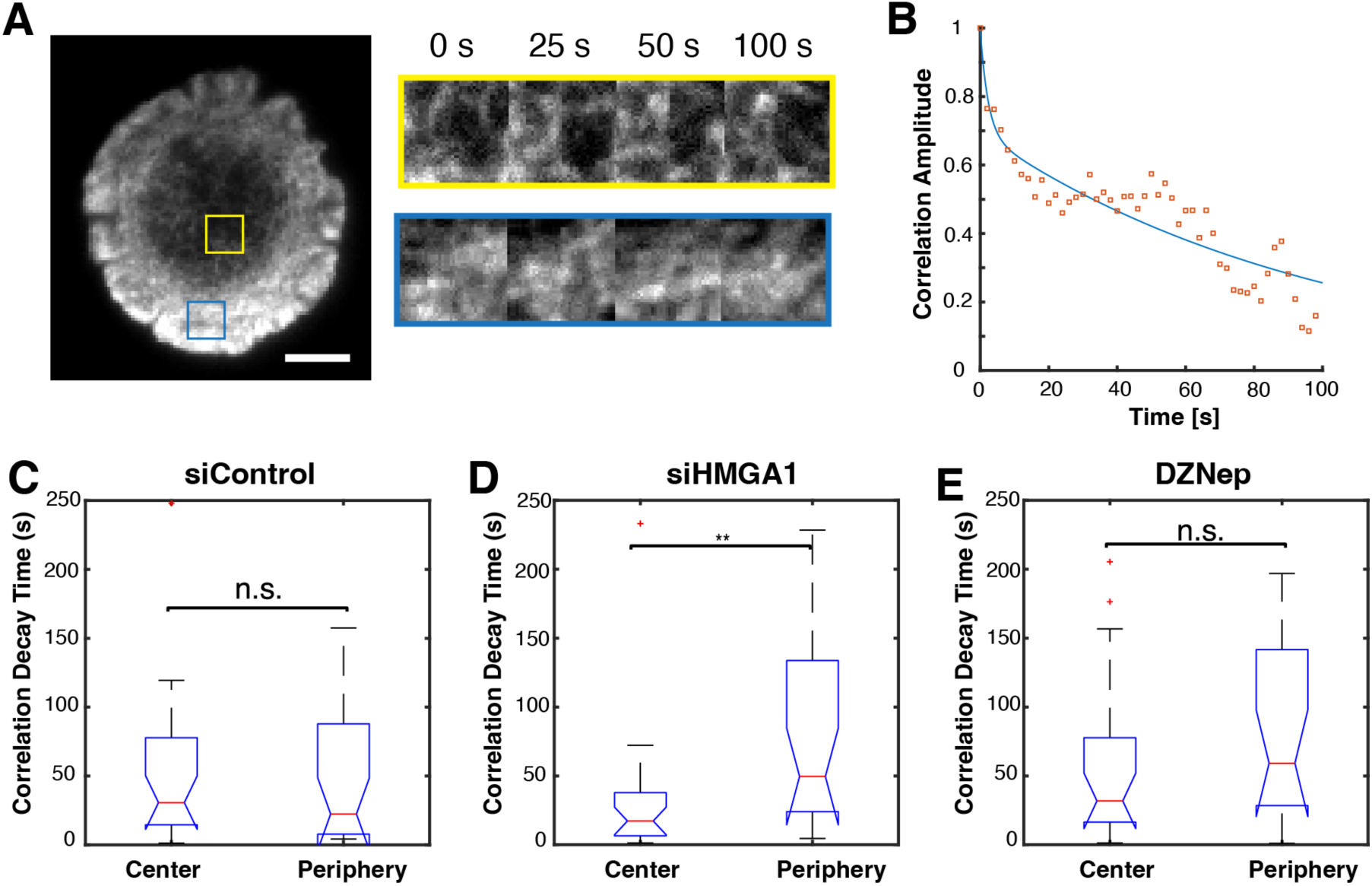
Chromatin compaction affects actin dynamics at the immune synapse. A) TIRF snapshot of activated tdTomato-F-tractin expressing Jurkat T cell and montage showing the actin structures at different times for a central (yellow) and peripheral (blue) region. B) Plot showing the correlation amplitude decay over time (orange squares) and the resulting fitted function (blue line) for the peripheral region of a sample siControl cell. Plots of the correlation decay time at the center and periphery regions for siControl (n=14 cells) (C), siHMGA1 (*p*<0.01 Wilcoxon rank sum test, *n*=13 cells) (D), and DZNep treated cells (*n*=13 cells) (E). Scale bar is 5 μm.

### Chromatin compaction modulates microtubule tip dynamics

Previous studies have shown that actin can modulate the speed and directionality of microtubule (MT) growth and tip motion during T cell activation^36, 38^. These result from direct interactions between actin filaments and growing microtubule tips, or by steric hindrance from the actin meshwork. We next investigated whether changes in actin network morphology induced by chromatin compaction influenced microtubule dynamics as well. To study MT tip dynamics, EGFP-EB3 (which marks newly growing MT plus-ends) expressing T cells were allowed to activate and spread for 5 minutes on anti-CD3 coated coverslips and imaged using TIRF for 5 minutes. Figure 5A shows the maximum intensity projection (color coded for time) corresponding to a representative time series of EB3. The EB3 comet-like particles were tracked using Utrack^39^ to obtain MT tip trajectories and quantify their dynamics.

**Figure 5.**
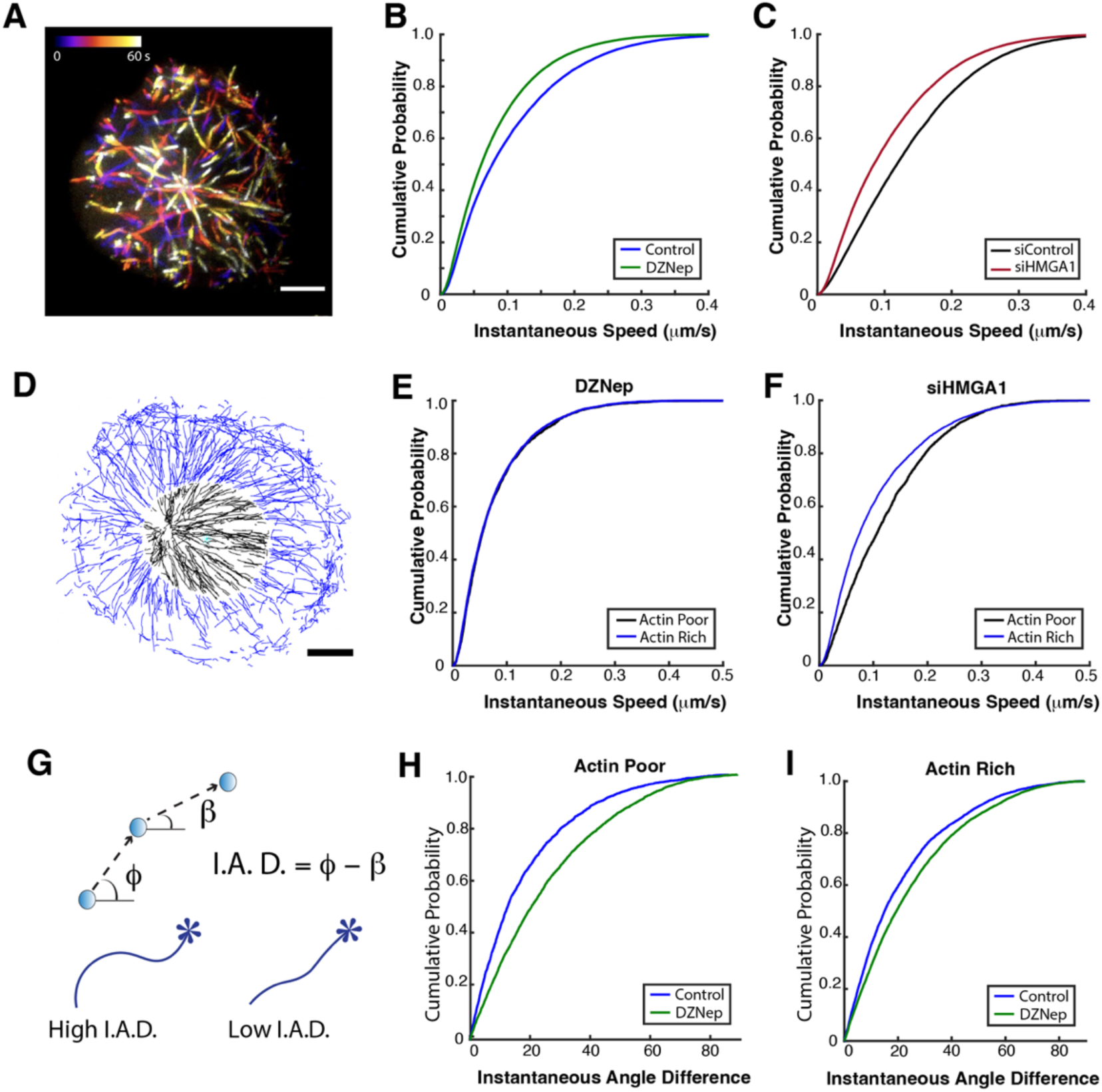
Chromatin compaction state modulates microtubule growth dynamics. A) Temporal color-coded maximum intensity projection of EGFP-EB3 expressing Jurkat T cell imaged with TIRF. B) Cumulative distribution of instantaneous speeds of EB3 for control and DZNep treated cells (*p*<0.001 Kolmogorov-Smirnov test, n=11 cells for both conditions). C) Cumulative distribution of instantaneous speeds of EB3 for control cells and HMGA1 KD cells (*p*<0.001 Kolmogorov-Smirnov test, *n*=15 and *n*=13 cells). D) Image showing the EB3 tracks obtained for a cell and their classification based on location - actin poor region in black and actin rich region in blue. Plots comparing EB3 speeds for actin rich region (peripheral) and actin poor region (central) for DZNep treated cells (*p* > 0.1 Kolmogorov-Smirnov test, *n*=11 cells) (E) and HMGA1 knock down cells (*p*<0.001 Kolmogorov-Smirnov test, *n*=13 cells) (F). G) Diagram illustrating the Instantaneous Angle Difference (I.A.D.) as the difference of angles corresponding to subsequent displacements within a trajectory. Cumulative distribution of the instantaneous angle difference (change in directionality) for DZNep treated cells in the actin poor region (G) and actin rich region (H) (*p*<0.001 Kolmogorov-Smirnov test, *n*=11 cells for both plots). Scale bars are 5 μm.

We found that both an increase and a decrease in chromatin compaction state led to a decrease in overall EB3 speeds as compared to the control cases (Figure 5B - DZNep, Figure 5C - siHMGA1). To explore this in more detail, we classified the EB3 trajectories in different regions of the synapse based on the actin distribution and defined an actin-poor region (black) and an actin-rich region (blue), as shown in Figure 5D and described previously^36^. We found that DZNep-treated cells showed no difference in instantaneous speed distributions between the actin-rich and actin-poor regions (Figure 5E). This contrasted with our previous findings of decreased tip speeds in the actin-rich regions in the absence of perturbations. Interestingly, MT tip speeds showed the expected difference between actin-rich and actin-poor regions in siHMGA1 cells (Figure 5F) and in the controls for both DZNep and HMGA1 (Supplementary Figure 5A, B), with the actin-rich peripheral regions showing lower MT tip speeds.

MT growth rates are controlled by intrinsic factors (e.g. post-translational modification of tubulin) as well as interactions with other cytoskeletal networks (e.g. actin and vimentin)^40, 41^. Accordingly, in untreated or siHMGA1 cells, the slowdown of growing MTs in the actin-rich periphery is likely due to steric hindrance from the dense lamellar/lamellipodial actin meshwork. However, the slowdown of MT growth in both the cell periphery and the actin-poor interior in DZNep-treated cells cannot be explained by steric hindrance. Instead, the lack of a ramified synaptic actin network in DZNep-treated cells (given lower actin levels) likely leads to an overall decrease in the ability of the actin network to guide growing MT filaments. We thus hypothesized that MT filaments in DZNep-treated cells would show less directional persistence even in actin-poor regions compared to control cells. To test this, we computed the instantaneous angle difference as the difference in angles between two consecutive interframe displacements (Figure 5G). We found that DZNep treatment caused a significant increase in the instantaneous angle difference of microtubule tips in both actin poor (Figure 5G) and actin rich regions (Figure 5H) indicating frequent changes in EB3 directionality. siHMGA1 cells displayed no significant change in microtubule tip directionality in either actin poor or actin rich regions (Supplementary Figure 5C, D). Together, these results show that altering chromatin compaction leads to a reorganization of the synaptic actin network, which in turn affects MT dynamics, suggesting a reciprocal interaction between the nucleus and different components of the cytoskeleton.

### Chromatin modulation of cytoskeleton morphology is mediated by SUN proteins and Myosin

Given our observations of chromatin compaction-mediated cytoskeletal changes during T cell activation, we hypothesized that the LINC complex likely plays an important role in transmitting signals between the nucleus and cytoskeleton in response to changes in chromatin state. SUN proteins, central components of the LINC complex, are inner nuclear membrane proteins which connect the interior of the nucleus to the cytoskeleton. We used siRNA to knock down SUN1 (Supplementary Figure 6A, B) or SUN2 (Supplementary Figure 6 C, D). We found that SUN1 knockdown cells (siSUN1), displayed similar actin accumulation as siControl cells (Supplementary Figure 6E), while SUN2 knockdown cells had decreased in actin MFI at the synapse (Supplementary Figure 6F). Since we wanted to examine the role of the LINC complex in the chromatin compaction-mediated modulation of the cytoskeleton independent of its direct effect on the actin cytoskeleton, we focused on SUN1.

Confocal imaging of cells with double knockdown of HMGA1 and SUN1 (siHMGA1 + siSUN1) showed that the increase in actin accumulation at the synapse, characteristic of siHMGA1 cells, was significantly reduced in siHMGA1 + siSUN1 cells (Figure 6A, B). Furthermore, the enhancement of peripheral actin observed in siHMGA1 cells was reduced in siHMGA1 + siSUN1 double knockdown cells, which displayed an actin distribution resembling that of siControl cells (Figure 6C). Our observations indicate that SUN1 KD abrogates the effect of HMGA1 KD on actin organization at the immune synapse and points to its role in maintaining nuclear cytoskeletal interactions. The nuclear deformation ratios of the siHMGA1 + siSUN1 double knockdown cells were slightly higher than those of siHMGA1 cells (Figure 6D). Though the difference was not statistically significant, this nevertheless suggests that SUN1 loss may in part rescue the effect of HMGA1 KD on nuclear deformations. SUN1 KD also reversed the effects of DZNep on actin intensity at the synapse, and to some extent the center to periphery ratio (Supplementary Figure 6G, H). Together, our results suggest that the mechanical connection between the nucleus and cytoplasm mediated by SUN1 plays a key role in linking chromatin compaction to cytoskeletal morphology.

**Figure 6.**
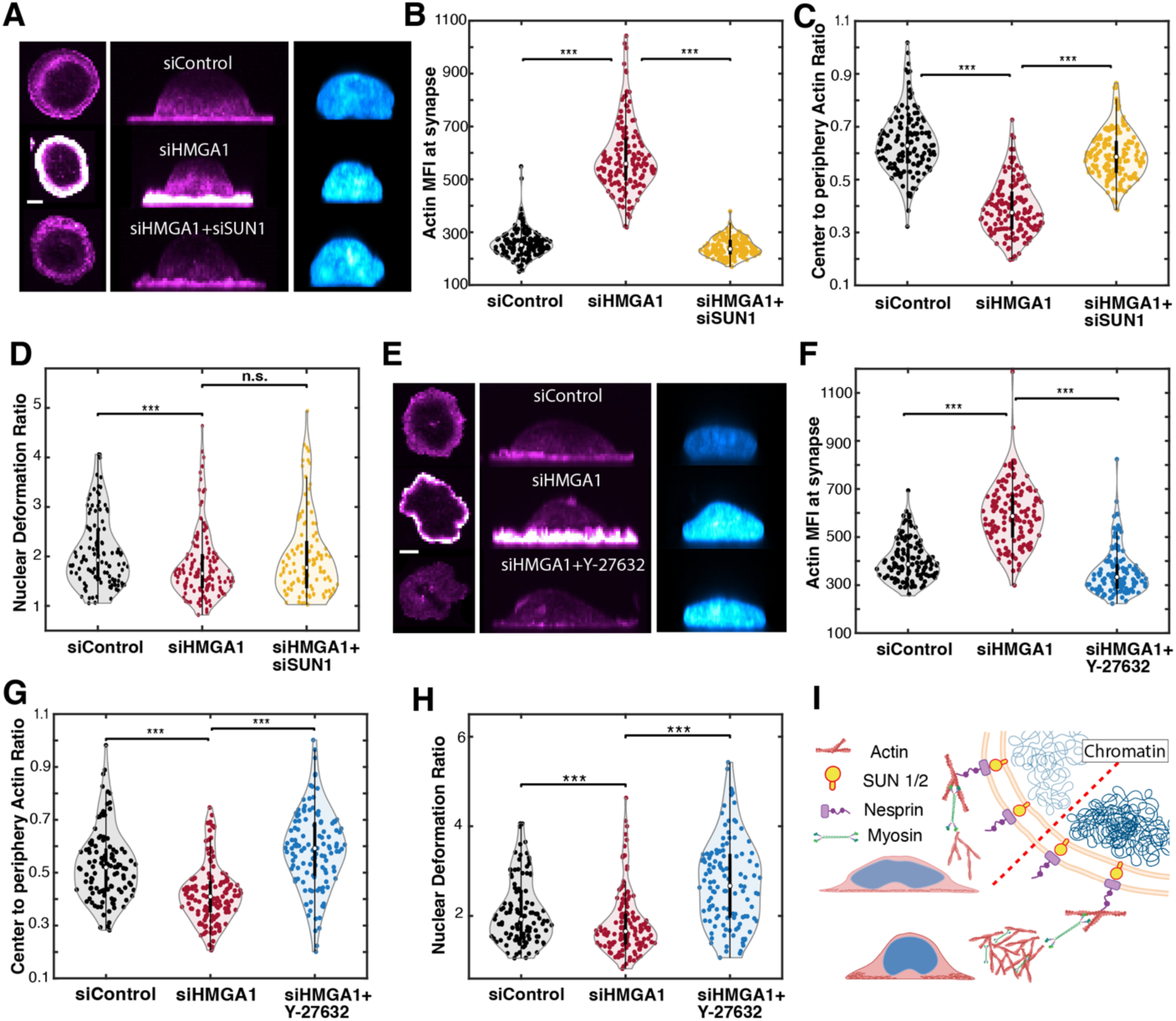
SUN1 and Myosin mediate bidirectional chromatin-actin cytoskeleton interactions. A) Synapse slice (left panels) and maximum intensity projections in *xz* (middle and right panels) of actin (magenta, labeled with phalloidin) and the nucleus (blue, labeled with Hoechst). Images are shown with the same intensity display range. B) Plots of F-actin MFI at the contact zone (synapse) for siControl, siHMGA1 and siHMGA1 + siSUN1 cells (siControl vs siHMGA1: *p*<0.001 Wilcoxon rank sum test, siHMGA1 vs siHMGA1+siSUN1: *p*<0.001 Wilcoxon rank sum test, *n*=135, n=140 and n=150 cells). C) Actin distribution measured by center to periphery ratio for siControl, siHMGA1 and siHMGA1+siSUN1 cells (siControl vs siHMGA1: p<0.001 Wilcoxon rank sum test, siHMGA1 vs siHMGA1 + siSUN1: *p*<0.001 Wilcoxon rank sum test, *n*=135, *n*=140 and n=150 cells). D) Nuclear deformation ratio measured in siControl, siHMGA1 and siHMGA1 + siUNC84a cells (siControl vs siHMGA1: *p*<0.001 Wilcoxon rank sum test, siHMGA1 vs siHMGA1 + siSUN1: *p*>0.1 Wilcoxon rank sum test, *n*=135, n=140 and *n*=150 cells). E) Synapse slice (leftmost panels) and maximum intensity projections in *xz* (middle and right panels) of actin (magenta, labeled with phalloidin) and the nucleus (blue, labeled with Hoechst) for siControl, siHMGA1 cells and siHMGA1 cells treated with Y27632 to inhibit myosin activity. Images are shown with the same intensity display range. F) Plots of actin MFI at the synapse measured for siControl cells, siHMGA1 cells and siHMGA1 cells treated with Y27632. G) Plots of actin center to periphery ratio for siControl cells, siHMGA1 cells and siHMGA1 cells treated with Y27632. H) Nuclear deformation ratio measured in siControl cells, siHMGA1 cells and siHMGA1 cells treated with Y27632. I) Representative schematic showing how chromatin compaction state may modulate the actin cytoskeleton via SUN1 and myosin. Increase in chromatin compaction induces higher actin accumulation in the periphery of activated cells and leads to reduced spread area and nuclear deformation. Decrease in chromatin compaction induces lower actin accumulation but leads to increased spread area and nuclear deformation. These changes require SUN1 and myosin. Scale bars are 5 μm.

Myosin has been identified to exert tension on nesprin^42^, a component of the LINC complex. In turn, the interaction between nesprin and SUN proteins has been reported to be indispensable for proper force transmission between the nucleus and the cytoskeleton^43^. To investigate the role of myosin, we used the ROCK-kinase inhibitor Y27632 to inhibit myosin phosphorylation and associated cell contractility. We treated siHMGA1 cells with Y27632 and used confocal microscopy to compare these cells with untreated siControl and siHMGA1 cells (Figure 6E). Y27632 treatment reversed the enhanced actin accumulation (Figure 6F) and the center to periphery ratio of actin at the synapse observed in siHMGA1 cells (Figure 6G). Furthermore, the inhibition of myosin activity also induced an increase in nuclear deformation of the cells, reversing the effect of HMGA1 KD (Figure 6H).

In summary, we find that an increase in chromatin compaction (nuclear stiffening) induces higher accumulation of actin in activated T cells. The increase in actin occurs mostly at the periphery of the cell and is correlated with reduced spread area and nuclear deformation. Decrease in chromatin compaction (nuclear softening) induces reduced actin accumulation but leads to increased spread area and nuclear deformation. Our results indicate that these alterations in cytoskeletal morphology as a function of chromatin compaction require both SUN proteins and myosin (Figure 6I).

### Chromatin compaction influences cellular mechanosensitivity

Immune cells encounter a variety of mechanical cues in vivo, such as stiffness of the antigen presenting surface of APCs, which can modulate their cytoskeletal organization and functional responses^44, 45^. How T cells adapt their nuclear and cytoskeletal dynamics to respond to mechanical cues such as stiffness, topography or external forces is an open question. Modulation of chromatin compaction by substrate stiffness has been reported for some cell types^19–21^. Conversely, manipulations of chromatin compaction can alter cellular response to stiffness changes^19, 21^. These studies point to a potential role for chromatin organization in regulating cellular mechanoresponse across cell types.

To test whether chromatin compaction modulates T cell mechanosensitivity, we fabricated polyacrylamide gels of tunable stiffness as described in our previous work^22, 46^. We examined the effect of chromatin decompaction by activating control and DZNep-treated cells on activating anti-CD3 coated polyacrylamide gels of 1 kPa and 12 kPa stiffness. These stiffness values span the range encountered by T cells^45–48^ and have been shown to modulate T cell activation and spreading. Cells were allowed to spread on activating gels, fixed and stained with phalloidin and Hoechst to visualize actin and the nucleus respectively (Figure 7A: control, Figure 7B: DZNep). Both control and DZNep-treated cells had larger spread areas on stiff (12 kPa) gels than on soft (1 kPa) gels (Figure 7C). The spread area of DZNep-treated cells was higher than that of control cells for both stiffnesses, consistent with our observations on glass. The nucleus deformation ratio was also higher for cells on stiffer gels, consistent with the expected correlation between cell spread area and nuclear deformation ratio (Figure 7D). For each stiffness, the nucleus deformation ratio was higher for DZNep-treated cells than for control cells, similar to glass. Finally, we found that cells activated on stiff gels displayed higher actin accumulation than cells on soft gels, for both control and DZNep conditions (Figure 7E). Actin accumulation was lower for DZNep-treated cells than for the control in each stiffness, indicating that nuclear decompaction (softening) leads to increased cell spreading, nuclear deformation and decreased actin accumulation on different stiffnesses (as shown schematically in Figure 7F).

**Figure 7.**
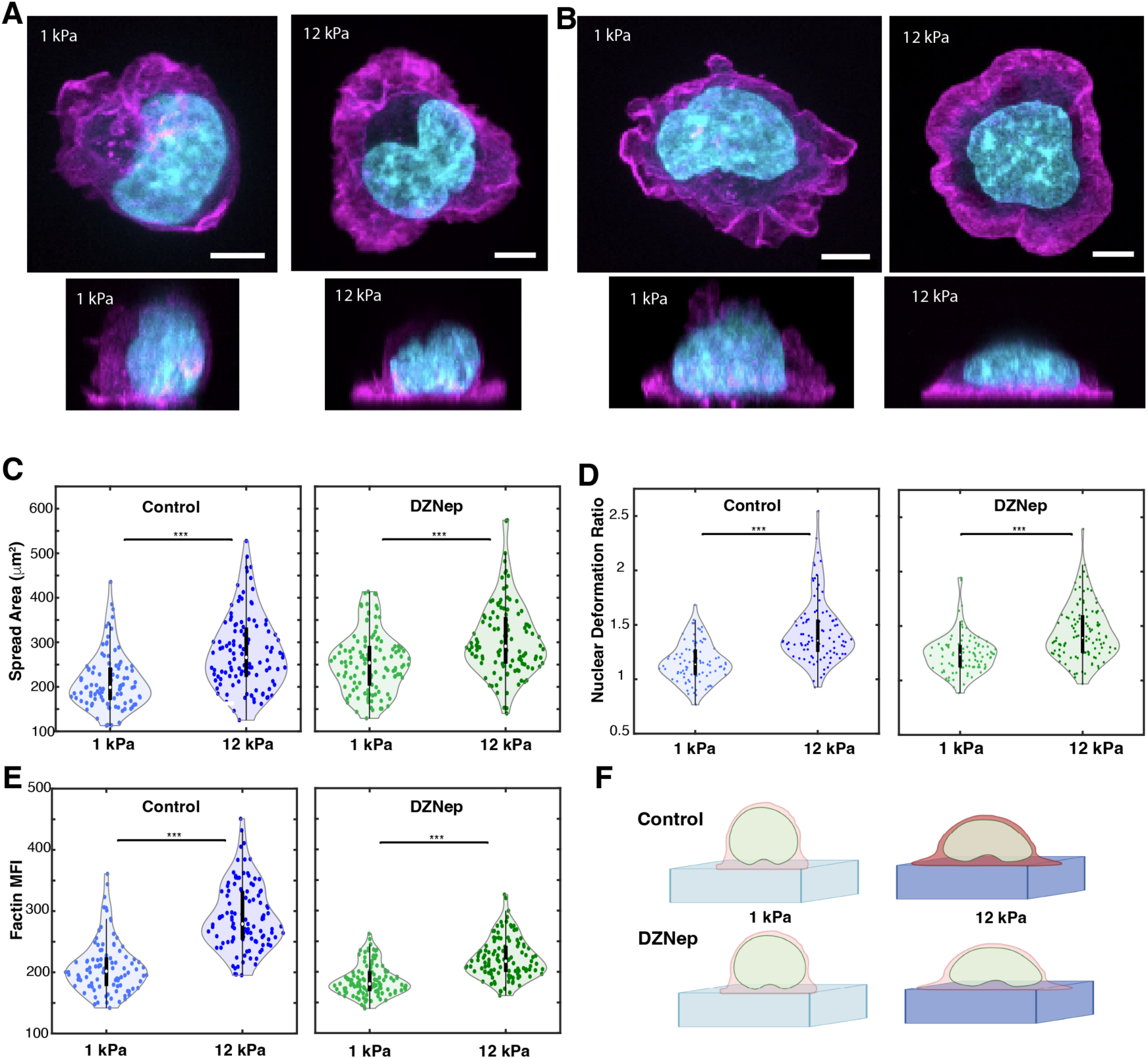
Reduced chromatin compaction influences cell response to stiffness. A) Maximum intensity projections in *xy* (top panels) and *xz* (bottom panels) of control Jurkat T cells activated on 1 kPa and 12 kPa gels. B) Maximum intensity projections in *xy* (top panels) and *xz* (bottom panels) of DZNep treated cells activated on 1 kPa and 12 kPa gels. Cells are for labeled for actin with phalloidin (magenta) and the nucleus with Hoechst (cyan) in A and B. C) Plots comparing the spread area of activated cells on 1 kPa and 12 kPa stiffness gels for control and DZNep treated cells (Control: *p*<0.001 Wilcoxon rank sum test, *n*=96 and 111 cells; DZNep: *p*<0.001 Wilcoxon rank sum test, *n*=117 and 118 cells). D) Nuclear deformation ratio for control and DZNep treated cells activated on gels of 1 kPa and 12 kPa stiffnesses (Control: *p*<0.001 Wilcoxon rank sum test, *n*=90 and 97 cells; DZNep: *p*<0.001 Wilcoxon rank sum test, *n*=94 and 111 cells). E) Plots showing actin accumulation for cells on 1 kPa and 12 kPa stiffness gels for control cells and cells treated with DZNep (Control: *p*<0.001 Wilcoxon rank sum test, *n*=96 and 111 cells; DZNep: *p*<0.001 Wilcoxon rank sum test, *n*=117 and 118 cells). F) Schematic of T cells activated on gels of indicated stiffness. Actin is depicted in red (with darker color representing higher actin accumulation). T cells display higher spread areas and higher F-actin accumulation in higher stiffness substrates. The stiffness-dependent increase of actin is diminished upon chromatin decondensation by DZNep treatment. Scale bars are 5 μm.

Given the effects of chromatin decompaction on the cellular response to stiffness, we next examined how increasing chromatin compaction would impact stiffness dependent changes in cell morphology. We activated siControl (Figure 8A) and siHMGA1 cells (Figure 8B) on gels of 1 kPa and 12 kPa stiffness. While control cells displayed higher spread area on stiffer substrates (Figure 8C), the spread area of siHMGA1 cells was not significantly different across the stiffnesses studied. On the other hand, the spread area of siHMGA1 cells was lower than control cells on 12 kPa gels, recapitulating the effect observed on glass. Despite the lack of difference in spread area for siHMGA1 cells, the nucleus deformation ratio was higher on the 12 kPa gels (Figure 8D) than on the 1 kPa gels for siControl and siHMGA1 cells. Finally, we quantified actin accumulation at the contact zone and found that the actin MFI was higher on 12 kPa gels than on 1 kPa gels for both siControl and siHMGA1 cells (Figure 8E). However, the stiffness-dependent difference in actin accumulation was smaller for the siHMGA1 cells. These observations are represented schematically in Figure 8F.

**Figure 8.**
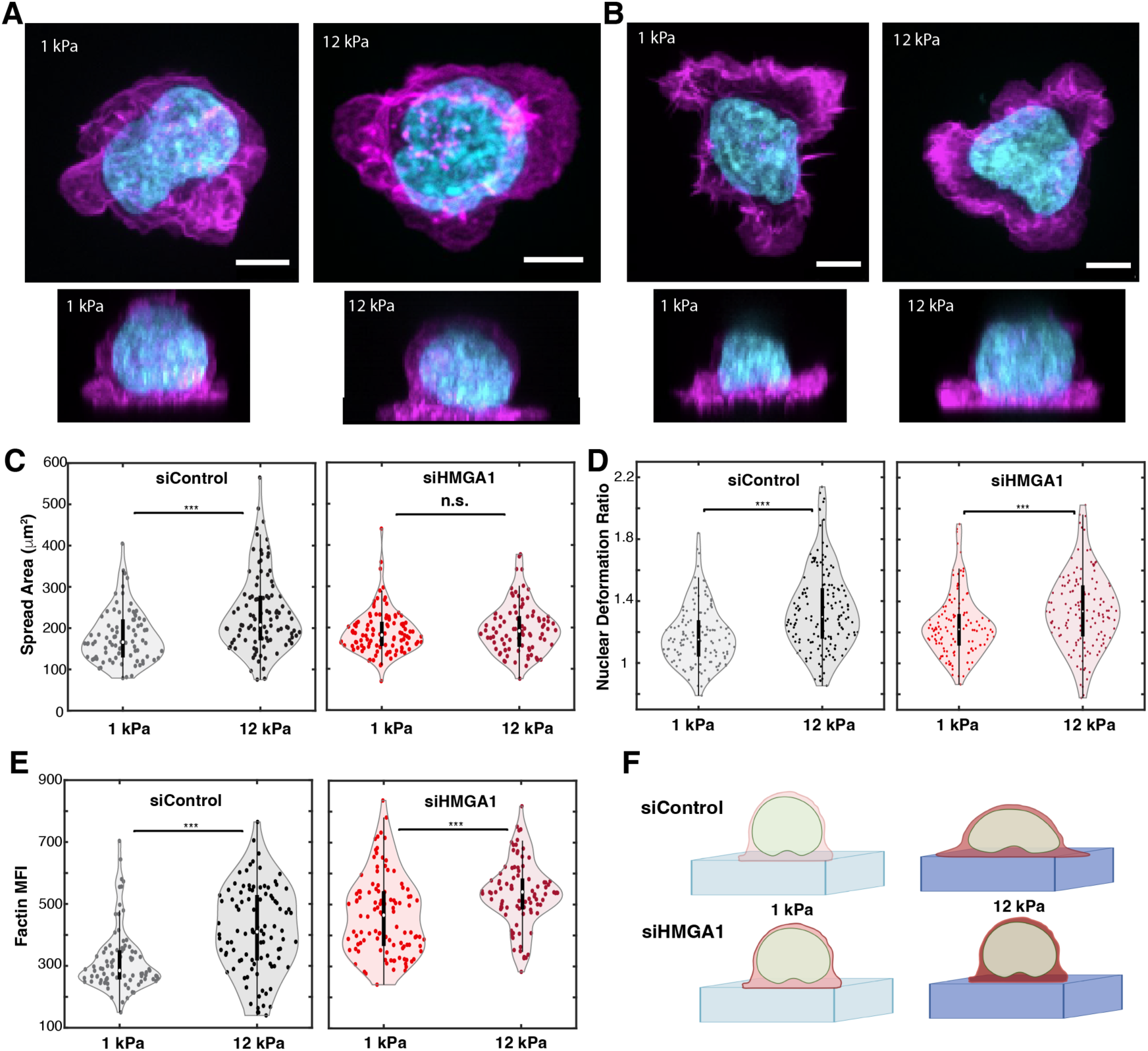
Increase in chromatin compaction modulates cell response to stiffness. A) Maximum intensity projections in *xy* and *xz* of siControl cells activated in 1 kPa and 12 kPa gels. B) Maximum intensity projections in *xy* and *xz* of siHMGA1 cells activated in 1 kPa and 12 kPa gels. Cells are for labeled for actin with phalloidin (magenta) and the nucleus with Hoechst (cyan) in A and B. C) Spread area of activated control siRNA and HMGA1 knock down cells on gels of 1 kPa and 12 kPa stiffness (siControl: *p*<0.001 Wilcoxon rank sum test, *n*=85 and 99 cells; siHMGA1: *p*>0.1 Wilcoxon rank sum test, *n*=108 and 92 cells). D) Comparison of nuclear deformation for siRNA control and HMGA1 knock down cells activated on 1 kPa and 12 kPa gels (siControl: *p*<0.001 Wilcoxon rank sum test, *n*=133 and 156 cells – siHMGA1: *p*>0.1 Wilcoxon rank sum test, *n*=129 and 139 cells). E) F-actin MFI at the synapse for cells activated on 1kPa and 12kPa gels (siControl: *p*<0.001 Wilcoxon rank sum test, n=85 and 99 cells – siHMGA1: *p*<0.001 Wilcoxon rank sum test, *n*=108 and 92 cells). F) Schematic of T cells activated on gels of indicated stiffness. Actin is depicted in red (with darker color representing higher actin accumulation). T cells display higher spread areas and higher F-actin accumulation in higher stiffness substrates. The stiffness-dependent increase in spread area is abrogated upon chromatin condensation increase by HMGA1 knock down. Scale bars are 5 μm.

A two-factor ANOVA analysis (with vehicle and inhibitor or siControl and siHMGA1 as one factor and the two stiffness values as a second factor) showed that DZNep treatment significantly attenuated the stiffness-dependent increase in actin MFI (F value for drug:stiffness interaction = 35.3, *p* = 5.7×10^-9^) while it had no effect on stiffness-dependent cell spreading (F value for drug:stiffness interaction = 0.75, *p* = 0.4). In contrast, HMGA1 knockdown significantly attenuated the stiffness-dependent increase in cell spread areas (F = 12.4, *p* = 0.005), but it did not have a significant effect in actin accumulation at the synapse (F = 0.85, *p =* 0.35). This suggests a dissociation of the effect of chromatin compaction and decompaction on different aspects of mechanosensitivity during immune synapse formation.

Nucleus deformation during T cell spreading is likely a result of cytoskeletal forces at the distal cell edge^10^ that are transmitted to the nucleus. We next turned to traction force microscopy to get a measure of the exerted forces and how changes in chromatin compaction affects these forces. siControl or siHMGA1 T cells were allowed to spread on anti-CD3 coated polyacrylamide substrates embedded with fluorescent 100 nm diameter beads (1kPa stiffness) and the resulting tractions were computed using bead displacements^46^. We found that despite the large variability in the overall total force exerted, siControl and siHMGA1 cells exerted similar levels of traction (Supplementary Figure 7A-C). Comparatively, vehicle-treated or DZNep-treated cells also exerted similar levels of traction (Supplementary Figure 7D-F).

## Discussion

In this study, we used confocal microscopy and quantitative image analysis to characterize nuclear morphology and chromatin organization during T cell activation. We found that the nucleus, in both primary and immortalized T cells, is deformed and compressed as the T cell spreads, and that these deformations are accompanied by an increase in chromatin compaction, which may contribute to the observed increase in T cell stiffness during activation^49^. To determine the role of chromatin compaction in T cell spreading and establishment of the immune synapse, we used small molecule inhibitors and RNA interference to alter chromatin compaction states. We found that decreasing chromatin compaction increased cell spread area and nuclear deformation but reduced F-actin accumulation and peripheral actin enrichment. Conversely, increased chromatin compaction led to reduced spread area and nuclear deformation. This was accompanied by enhanced levels of peripheral F-actin at the immune synapse. Our results imply a reciprocal interaction between chromatin compaction and the organization of the actin cytoskeleton at the immune synapse.

The changes in cytoskeletal network morphology at the cell-substrate interface induced by changes in chromatin compaction could result from several complementary mechanisms. It is possible that these changes in actin at the synapse are a result of feedback mechanisms^50^ between nuclear shape and cytoskeletal organization and contractility. One possibility is that, when the nuclear height reaches a threshold due to compressive forces, it results in stimulation of actin polymerization and actomyosin contractility to counteract these forces. Chromatin decompaction by DZNep likely enables the nucleus to accommodate these compressive forces without the need to engage stretching of the nuclear membrane and subsequent increase in actin polymerization. Conversely, increased chromatin compaction by knockdown of HMGA1 lowers the threshold height required for engagement of this feedback, leading to enhanced actin accumulation at the synapse. This view is also consistent with our observations that the total traction forces generated by the cells is largely unchanged despite alterations in chromatin compaction, suggesting that these cells have maximal force generation capability that limits how much they can accommodate or compress a stiffer or softer nucleus.

The increased actin enrichment observed with enhanced chromatin compaction in siHMGA1 cells requires the LINC complex. SUN1 knockdown abrogated the altered actin distribution and accumulation in siHMGA1 cells, indicating a LINC-mediated interplay between the nucleus and cytoskeleton. SUN proteins have also been shown to regulate the level of histone methylation in the nucleus^51^ and SUN1/2 KD leads to a reduction in H3K27 trimethylation and a redistribution of histones and chromatin decompaction. SUN1-mediated decompaction likely counteracts the effect of HMGA1 knockdown, restoring chromatin compaction (and hence mechanical properties) to near control levels. Additionally, the LINC complex is subject to forces produced by myosin motors, which may result in reorganization of chromatin^39^. Inhibition of Rho-kinase activity and myosin contractility abolished the increase in synaptic F-actin induced by chromatin compaction. Thus, it is possible that the tension produced by myosin is required for proper communication between the nucleus and the cytoskeleton. Consistent with this view, LINC complex-mediated communication between the nucleus and the cytoskeleton has been reported for other cell types. B cell nuclear morphology adapts in a cytoskeleton-dependent manner, requiring the LINC complex and playing a critical role in organized immune synapse formation^52^. Endothelial cell adaptation to external forces requires LINC complex-mediated cytoskeletal reconfiguration^53^ and the interaction between nesprins and SUN proteins (an intact LINC complex) is critical for force transmission between the nucleus and the cytoskeleton^43^.

It is well established that T cell spreading and activation is sensitive to substrate stiffness^2, 45–47^. In this study, we found that manipulation of chromatin compaction altered the mechanosensitivity of T cells. Increased cell spreading on stiff surfaces was accompanied by higher actin accumulation at the periphery. We observed that DZNep treatment and associated chromatin decompaction reduced the stiffness-dependent increase in actin, yet the spread area remained responsive to increasing stiffness. This suggests that nuclear softening due to chromatin decompaction may facilitate cell spreading at higher stiffnesses without a concomitant increase in actin polymerization. Conversely, increasing chromatin compaction abrogates the stiffness dependent increase in spread area, while maintaining the increase in F-actin levels as a function of stiffness. Thus, chromatin compaction and nuclear deformations play a key role in T cell mechanosensitivity.

Nuclear deformation of activated T cells has been reported for various geometries of antigen presenting surfaces^45, 46, 54–56^. Activation induced nuclear deformation accompanied by signaling is important for gene expression in adherent cells^57^. Nuclear deformation and associated stretching of the nuclear membrane induce opening of nuclear pore complexes and ion channels facilitating nuclear import and calcium signaling, essential in multiple cellular functions^12^. These results, together with our observations, suggest that nuclear deformations play an important physiological role during T cell activation rather than merely being a morphological consequence of activation.

Overall, our results are broadly consistent with the nuclear drop model for nuclear morphological changes during cell spreading ^9, 10, 58^. The in-plane stress generated by actin polymerization during cell spreading is transferred as a normal stress on the nucleus, leading to its deformation. In cells that have robust Lamin A/C expression, the nucleus deforms at constant volume until the nuclear membrane folds are lost. Since the nuclear lamina cannot accommodate strain, the cell stops spreading. However, given the limited expression of Lamin A/C in T cells, the bulk modulus of the nucleus plays a larger role. Several prior studies have implicated histone acetylation^9, 32, 57, 59^ and methylation^13, 18^ and nucleosome-nucleosome interactions in regulating overall chromatin compaction and thereby stiffness, especially at small strain regimes. Alterations of chromatin compaction, which determines the bulk modulus^13^, may thus have a significant effect on T cell spreading. In support of this model, we find that inhibition of myosin contractility by Y27632, which is known to enhance cell spreading^60, 61^ leads to higher deformation of nuclei under conditions of higher compaction. Our findings highlight the importance of chromatin modifications and epigenetic states in determining the spreading dynamics of cells, especially in those with limited Lamin A/C expression (such as T cells).

The nucleus not only houses and protects the genome, but also behaves as a mechanical element, both limiting morphological changes of cells and adapting to altered extracellular geometries^12, 62^. Nuclear deformation is a key component of the optimal regulation of cell function. Overall, our findings highlight the crucial role of chromatin conformation and associated nuclear mechanics in shaping T cell morphology and function during activation. Further studies are needed to fully elucidate the molecular mechanisms underlying this mechanical interplay between the nucleus and the cytoskeleton during T cell activation.

## Methods

### T cell activation

Coverslips attached to eight-well Labtek chambers were incubated in poly-l-lysine (PLL) at 0.01% W/V (Sigma Aldrich, St. Louis, MO) for 10 min. PLL was aspirated and the slide was left to dry for 1 h at 37°C. T cell activating antibody coating was performed by incubating the slides in a 10 μg/ml solution of anti-CD3 antibody (Hit-3a, eBiosciences, San Diego, CA) for 2 h at 37°C or overnight at 4°C. Excess anti-CD3 was removed by washing with L-15 imaging media right before the experiment.

### Cell culture and transient transfections

E6-1 Jurkat cells were cultured in RPMI 1640 supplemented with 10% fetal bovine serum (FBS) and 1% Penn-Strep antibiotics. For transient transfections with plasmids or siRNA, we used the Neon (ThermoFisher Sci.) electroporation system 2 days before the experiment. The protocol was as follows: The protocol was as follows: 2×10^5^ cells were resuspended in 10 μL R-buffer containing 0.5–2 μg plasmid DNA, electroporated with three pulses at 1325 V for 10 ms, and transferred into 550 μL pre-warmed RPMI 1640 with 10% FBS. For siRNA: 2×10^5^ cells were resuspended in 10 μL R-buffer containing 11 pmol siRNA, electroporated under the same conditions, and immediately transferred into 550 μL pre-warmed RPMI 1640 with 10% FBS, yielding a final concentration of ∼20 nM siRNA per well. The cells were exposed to three pulses of amplitude 1325 V and duration 10 ms in the electroporator. Cells were then transferred to 500 μl of RPMI 1640 supplemented with 10% FBS and kept in the incubator at 37°C. The tdTomato-F-tractin plasmid was a gift from John A. Hammer and the EGFP-EB3 plasmid was a gift from Robert Fisher. For knockdowns, we purchased Silencer Select siRNA from Invitrogen [HMGA1 (#4392420, ID: s6667), SUN1 (#4392420, ID: s23629), SUN2 (#4392420, ID: s24466)]. As a negative control, we used the Silencer Select Negative Control No. 1(#4390843).

### Inhibitor experiments

DZNep was purchased from EMD Millipore (#506069). Cells were incubated in 25 μM DZNep for 16-20 hours. For imaging of live and fixed cells, DZNep was also added to the wells at the same concentration used for incubation. Since DZNep is supplied dissolved in water, we added an equivalent volume of water (1:500 dilution) for the corresponding control condition. Y27632 (Calbiochem, Millipore-Sigma, Darmstadt, Germany) was used at 100 μM. For vehicle control experiments, cells were incubated for 5 min in a DMSO and L-15 solution at 0.01% concentration and the same concentration was kept in the imaging chamber.

### Immunofluorescence

Cells were activated on anti-CD3–coated coverslips, fixed after 7 min of activation using 4% paraformaldehyde for 10 min at room temperature, and then washed thoroughly with 1X PBS. Cells were then permeabilized for 5 -10 min with a 0.15% Triton X solution and blocked with BSA (0.02 g/ml) and glycine (0.3 M) in 1× PBS solution for 1 h at room temperature. For F-actin labeling, Acti-stain 670 phalloidin (Cytoskeleton, Denver CO) was used. Nucleus staining was performed using Hoechst 33342 from Invitrogen (#H3569) for fixed cells or with SiR-DNA from SpiroChrome (CY-SC007) following the manufacturer instructions. For incubation periods and concentrations of primary and secondary antibodies, we followed the manufacturers’ recommendations. For validation of protein knockdowns, immunostaining was performed using primary antibodies from Invitrogen; HMGA1 (PA5-78007), SUN1 (PA5-52564) and SUN2 (PA5-95741). Evaluation of chromatin condensation was performed with immunostaining of H3K27me3 with a primary antibody from Invitrogen (MA5-11198) and to measure early activation signaling a primary antibody for phosphorylated Zap70 from Cell Signaling Technology (#2701) was used. For secondary antibodies, we used Alexa Fluor Goat anti-Rabbit IgG (H+L) conjugates purchased from Invitrogen (#A-11008, # A-11012, # A-21070). Laser power and camera gains were set to be consistent across replicates and conditions to facilitate quantitative comparison.

### Polyacrylamide gel fabrication

Polyacrylamide hydrogels (stiffness 1 kPa and 12 kPa) were purchased from Matrigen (Softview 35/20 Glass #SV3520). The hydrogel surface contains quinone groups which form covalent bonds with proteins to enable functionalization. The gels were hydrated in 1xDPBS for 30 minutes before use, followed by extensive washing with 1xDPBS. Washed gels were coated with 10 µg/mL NeutrAvidin (Thermo Scientific #31000) at 4°C overnight, followed by washing with 1xDPBS. NeutrAvidin-coated gels were further incubated with 10 µg/mL biotinylated anti-human CD3 (BioLegend #300304) at 37°C for 2 hours and washed with warm Leibovitz’s L-15 imaging medium (Gibco #21083027). Jurkat T cells for the indicated conditions were centrifuged at 300g for 5 minutes, resuspended in L-15 imaging medium and added to coated polyacrylamide gels. The cells were allowed to adhere to the gel surface at 37°C and fixed at 7 minutes after addition with 3% paraformaldehyde for 10 minutes at room temperature in the dark, followed by extensive washing with 1xDPBS.

### Microscopy

Imaging was performed on an inverted microscope (Nikon Ti-E PFS, Melville NY) equipped with a laser TIRF illuminator and a CSU-W1 spinning-disk confocal unit (Yokogawa, Tokyo, Japan) and solid-state lasers of wavelength 491, 561 and 640 nm. A 100x 1.4 NA Silicone objective lens was used for confocal imaging, and a 100x 1.49 NA lens was used for TIRF imaging. Images were captured with a Prime BSI camera (Photometrics, Tucson AZ). Live imaging of cells was performed in L-15 medium medium with 5% FBS. Samples were equilibrated to 37°C in a stage-top Okolab Incubator (Okolab S. R. L., Pozzuoli, NA, Italy). For live confocal imaging, acquisition was initiated after cell addition and slices covering 15-20 um in z were captured with a z-spacing of 0.3 μm. Imaging volumes were acquired every 1 minute. For live TIRF imaging (of actin and EB3 dynamics), images were acquired every 2 s in two phases 0-5 min and 5-10 min after adding cells to activated surfaces. The data from 5-10 min of imaging was used for analysis. Imaging protocols were implemented using Nikon Elements. Images were cropped and dark image subtraction was performed in Fiji before further analysis using custom MATLAB scripts.

### Traction force microscopy

Cells were added to PA hydrogels and spinning-disk confocal images of SiR-DNA and the bead field were captured every 30 seconds with 488 nm and 640 nm laser lines. Images were captured 1.2 μm below and 10 μm above the gel surface with a spacing of 0.3 μm. Cells were detached from the substrate using 2 mL each of 0.25% trypsin-EDTA and 1x DPBS. A final image of the fluorescent bead field was acquired after detachment of the cells to be used as a reference frame.

### Image analysis

#### Nuclear segmentation

Individual cell images were cropped in Fiji and subsequent analysis was carried out in MATLAB (Mathworks, Natick MA). Each slice of the nuclear volume was smoothed with a 2D bilateral filter to reduce noise while preserving edges. The volume was resized to be isotropic, and background subtraction was performed. The *xy* maximum intensity projection was segmented using k-means clustering (k=2) followed by hole-filling along with morphological dilation and erosion operations. Only regions within the MIP mask were considered in subsequent steps. The broadest plane of the nucleus, identified as the plane with the highest total signal, was segmented analogously but omitting dilation so to avoid edge overexpansion.

The bottom and top planes of the nucleus were determined with a gradient-based boundary detection approach. First, axial intensity gradients were computed for each voxel above an intensity threshold. For Jurkat cells, this threshold was set to the 5th percentile of all values within the mask of the broadest slice of the nucleus, while for CTLs, due to differences in the internal intensity distribution, the threshold was set to half the median value within the mask. These axial gradients, influenced by both the distribution of fluorescence within the nucleus and by the point spread function of the imaging system, are generally positive just below the nuclear boundary and negative just above it. Accordingly, the bottom plane was conceptualized as the first axial slice for which intensity gradients were no longer consistently positive while the top was the first slice featuring heavily negative intensity gradients.

Concretely, let I denote a nuclear volume. 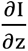 (**p**, z) is estimated for all x-y pixels **p** and planes z using finite central differences. Let P(z) denote the proportion of pixels in plane z above the intensity threshold for which the quantity 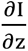 (**p**, z) > 0 and I(**p**, z) > *ε*, a small threshold (set to 10). Noting that P(z) ≈ 1 directly below the nucleus and that *P*(*z*) ≈ 0 directly above it, the axial range of the nucleus can be estimated as the set of planes *z*^∗^ = {z | a ≥ P(z) ≥ b} for some constants a and b. The constants a = 0.85 and b = 0.05 were manually selected and held consistent across experiments.

To segment the full nucleus, we first generated per-slice binary masks. For each slice between the computed bottom and top planes, an initial mask including all pixels above a given intensity threshold was generated. For Jurkat cells, this threshold was set to the 10^th^ percentile in intensity within the broadest slice; for CTLs, the threshold was instead set to half the median within this slice. However, signal from structures in nearby planes can lead to erroneous inclusions in the initial masks. To address this, we computed the normalized axial intensity gradient at each pixel. Pixels exhibiting both a high gradient magnitude and a consistent sign across neighboring slices— indicative of contributions from structures in nearby planes—were excluded from the mask. The resulting mask was filled in and refined morphologically by dilation and erosion operations. Finally, after reconstructing the full mask by stacking all planes, we applied 3D morphological closing and opening with a spherical structuring element to smooth boundaries and promote continuity across slices.

### 3D nuclear reconstructions and curvature computations

A 3D triangular mesh of the nuclear surface was generated using the MATLAB toolbox iso2mesh, and the mesh was subjected to 10 iterations of smoothing with the Humphrey’s Classes algorithm^63^ to refine the surface. The mesh was rendered using iso2mesh and the open-source MATLAB toolbox GIBBON.

Principal curvatures were computed over each face of the mesh to provide a local characterization of the nuclear surface^64^. Faces on the surface were classified as convex if both principal curvatures (*k*_1_and *k*_2_) were positive, hyperboloid if *k*_1_ > 0 and *k*_2_ < 0, and concave if *k*_1_ < 0 and *k*_2_ < 0. Following previous work^65^, the invagination ratio was computed as the ratio of the non-convex (hyperboloid and concave) surface area to the total surface area.

#### Nuclear deformation ratio and H3K27me3 Mean Fluorescence Intensity (MFI) calculations

The nuclear deformation ratio, a measure of nuclear flattening, was defined as the ratio of the nucleus’s lateral extent to its height. Lateral extent was computed as the average of the major and minor axis lengths of an ellipse sharing the same normalized second central moments as the projected nuclear region, as determined using *regionprops* in MATLAB. Nuclear height was defined as the difference between the maximum and minimum z-values among all faces on the mesh. The H3K27me3 MFI was computed by averaging background-subtracted voxel intensities from the H3K27me3 channel within the 3D nuclear mask.

#### Actin auto-correlation analysis

To characterize actin dynamics two square regions of interest (ROI) measuring 2.5 microns were selected at both the center and the peripheral areas of the cell (four per cell). The fluorescence signal within the ROI (Figure 4A) was registered at each timepoint, and the temporal autocorrelation was calculated by comparing the intensity value of the initial frame against itself and every subsequent frame. To quantify actin dynamics, the correlation amplitude time-course was fitted with a double-exponential curve of the form *C*(*t*) = *αe*^−*bt*^ + *ce*^−*dt*^. The value of α was constrained to a range close to 1, corresponding to the correlation at time zero (short times), with b the correlation decay rate at short time while c and d are the correlation amplitude and correlation rate at later times, respectively.

#### Traction force analysis

For 2D traction force analysis, confocal images of deformed bead volumes beneath the cells were aligned to corresponding undeformed reference images using a Fourier transform-based alignment method to achieve sub-pixel accuracy^66^. An *xy* MIP of the aligned bead images was generated to analyze bead displacements on the top surface of the gel. Bead displacements between the deformed and reference MIP images were calculated using high-resolution optical flow implemented in MATLAB. These displacement fields were then used to compute traction stress fields through using regularized Fourier transform traction cytometry (FTTC)^67^. The total force was calculated by integrating the traction forces over the footprint of the cell as imaged by brightfield microscopy.

### Statistical tests

MATLAB was used for statistical analysis and graphic representations of data. Non-parametric tests (two-tailed Wilcoxon rank-sum test with MATLAB function *ranksum* or two-sided Kolmogorov-Smirnov test with MATLAB function *kstest2*) were used for all statistical comparisons as indicated in the figure legends. Differences between values were considered statistically significant when *p* < 0.05 and non-significant (n.s.) for *p* > 0.05. The MATLAB function *anovan* was used for two-way ANOVA comparisons. Details of experimental conditions and numbers of observations are provided in the figure legends. All quantitative data were collected from 2-3 experimental replicates. All fixed cell results show data from one representative replicate in the figure. For all live cell data, the analysis was done with data pooled from all experiments.

## Acknowledgements

A.U. acknowledges support from the NIH (GM145313) and the NSF (12132922). I. R-S would like to acknowledge support from the Fulbright-Colciencias scholarship. We would like to thank Z. Xiao and L. Li (UMD) for providing primary CTLs. We acknowledge Z. Zimmerberg’s help with data analysis.

## Author contributions

I.R.-S. and A.U. designed the experiments with input from other authors. I.R.-S., A.P. and M.C. performed the experiments. I.R.-S. and F.F. designed and implemented the analysis pipeline and analyzed the data. I.R.-S. and A.U. wrote the draft of the manuscript, with input and editing from F.F., A.P. and M.C. A.U. supervised the project.

## Data Availability

All the data is available upon reasonable request.

