## Supplementary figures for "Nuclear morphology and chromatin organization modulate T cell cytoskeletal remodeling and immune synapse formation"

**Supplementary Information**

Supplementary Movie 1a: Timelapse images of maximum intensity projections in *x-y* (synapse view) of a Jurkat T cell spreading on an activating coverslip coated with anti-CD3. The nucleus is labeled with SiR-DNA (cyan) and actin is labeled with td-Tomato-F-tractin (magenta).

Supplementary Movie 1b: Timelapse images of maximum intensity projections in *x-z* (side view) of a Jurkat T (same cell as in Movie 1a) spreading on an activating coverslip coated with anti-CD3. The nucleus is labeled with SiR-DNA (cyan) and actin is labeled with td-Tomato-F-tractin (magenta).

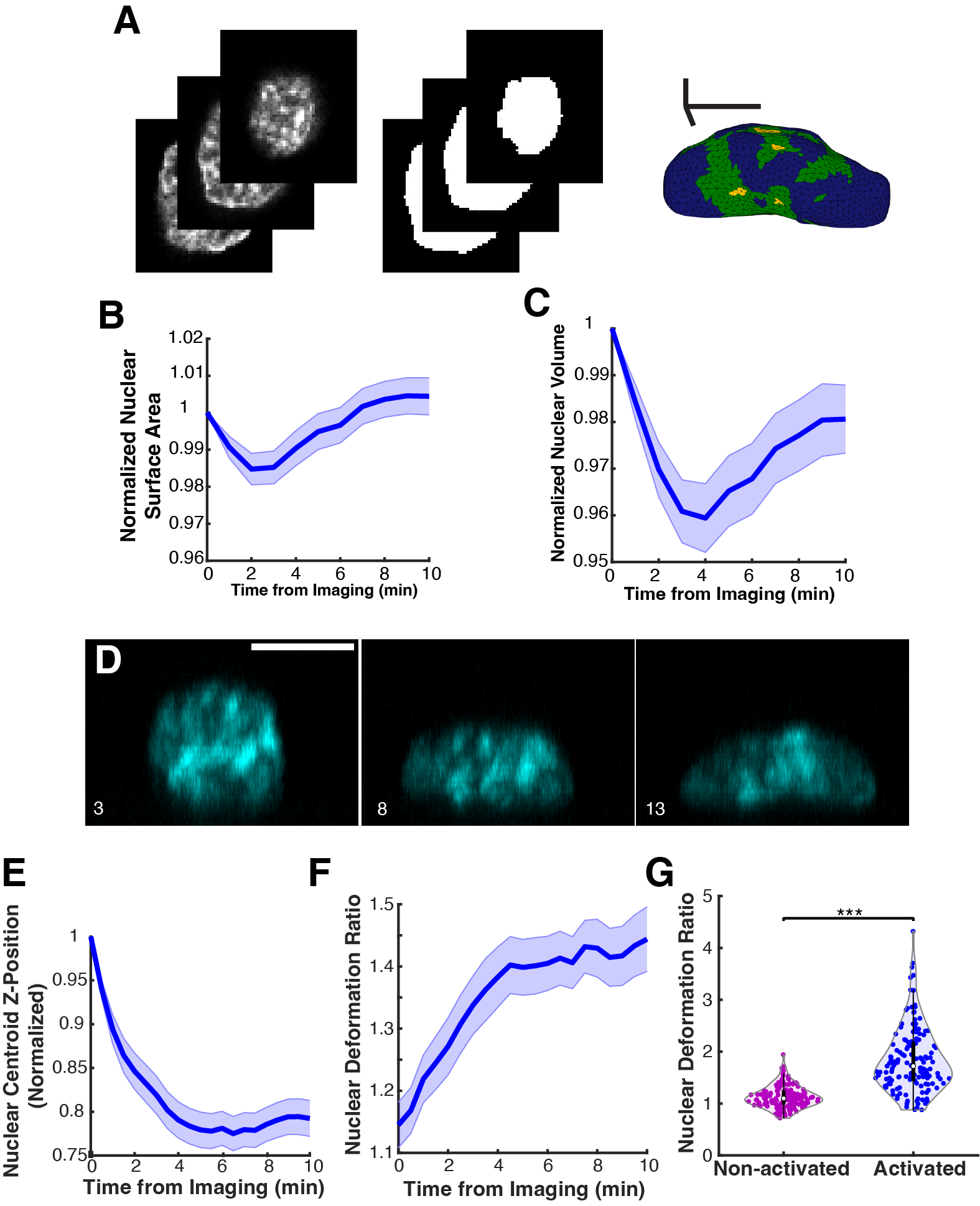

**Supplementary Figure 1. Workflow and analysis of nuclear shape deformation.**

A) Representative confocal images of the nucleus and their respective mask obtained after segmentation. The mesh model obtained from the mask is shown on the right. B) Normalized nuclear surface area as a function of time. C) Normalized nuclear volume as a function of time. Shaded portion of the curve is the standard error in both B and C. D) Maximum intensity projection in *xz* of the nuclear Hoechst signal for a primary CD8+ T cell activated on an anti-CD3 coated glass coverslip at the indicated times (3, 8, 13 minutes). E) Normalized nucleus centroid Z-position as a function of time for activated and spreading primary T cells. Shaded portion of the curve is the standard error (*n*=55 cells). F) Nuclear deformation ratio (ratio of height to average max radius) as a function of time for activated and spreading primary T cells. Shaded portion of the curve is the standard error (*n*=55 cells). G) Comparison of the deformation ratio for activated versus non-activated primary T cells (*p*<0.001 Wilcoxon rank sum test, *n* = 109 and 150 cells respectively). Scale bars are 5 μm.

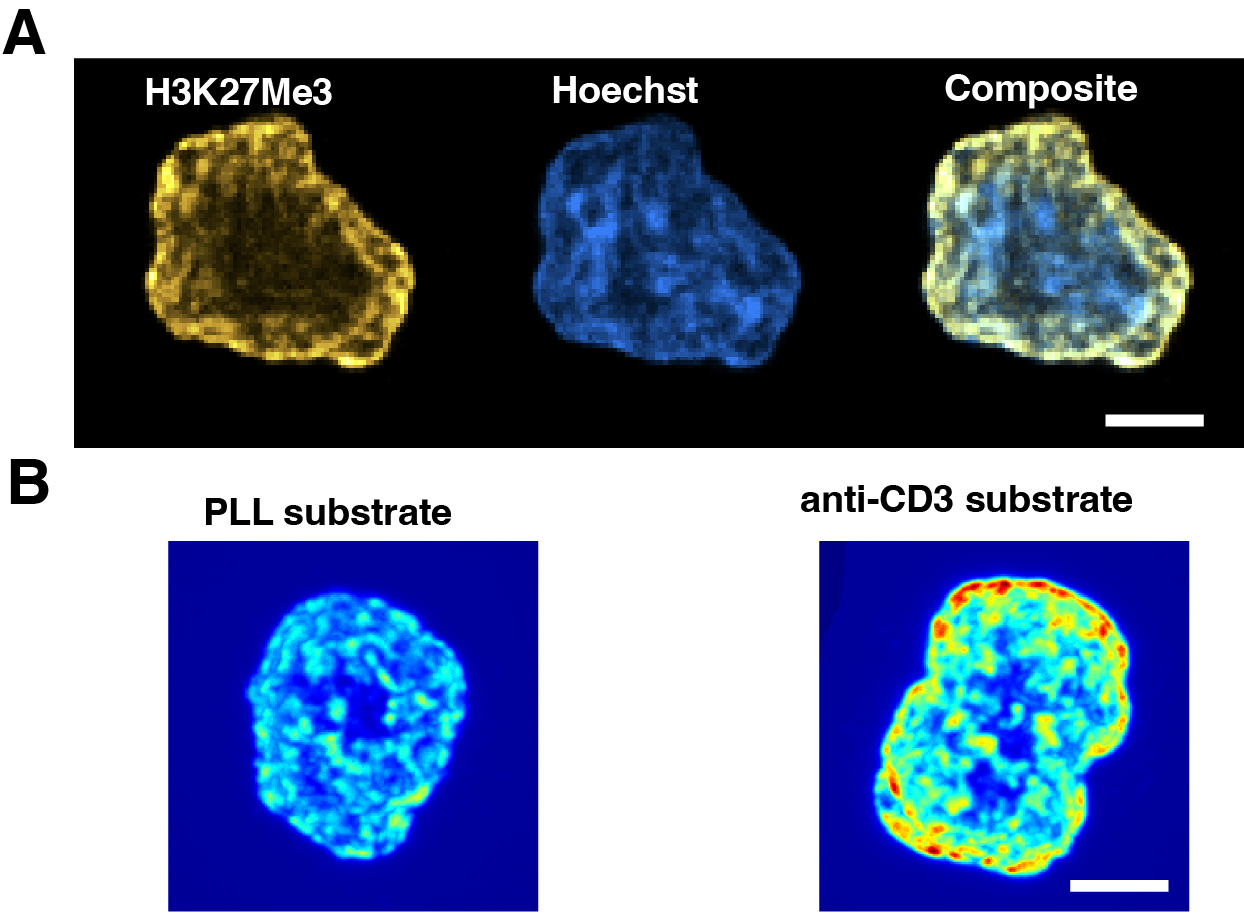

**Supplementary Figure 2: T cell activation enhances heterochromatin levels.** A) Confocal image (single slice) showing the distribution of the H3K27me3 marker (yellow) with reference to DNA stained with Hoechst (blue). B) Maximum intensity projections of H3K27me3 signal for cells on PLL (left) and anti-CD3 (right) coated coverslips. Scale bars are 5 μm.

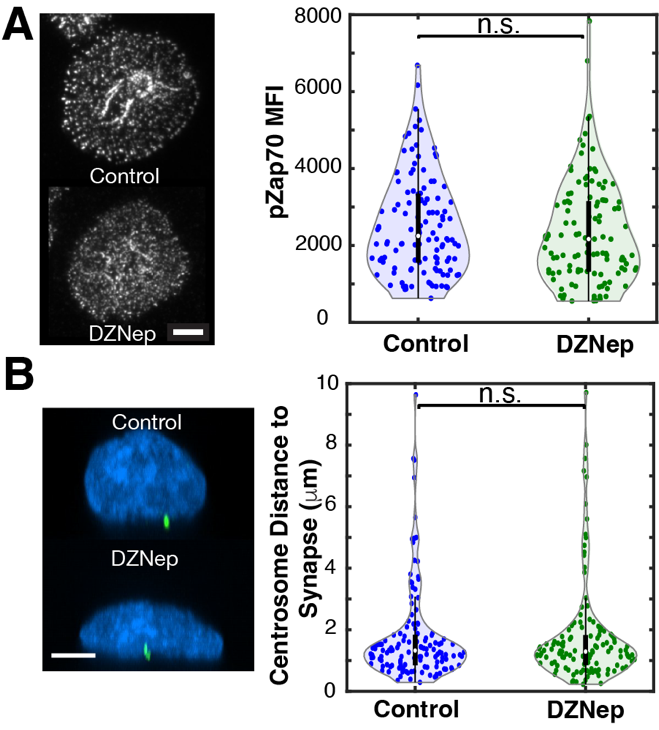

**Supplementary Figure 3: Chromatin decompaction does not affect early T cell signaling.**

A) Left panel: Representative TIRF images (at the synapse plane) of phosphorylated Zap70 in control and DZNep treated and activated Jurkat T cells. Right panel: Plots of pZAP70 mean fluorescence intensity for cells in the two conditions. B) Left panel: Representative maximum intensity *xz* projections of EGFP-centrin (green) expressing cells fixed and stained with Hoechst (blue) treated with DZNep or with control media. Right panel: plots of centrosome distance to the synapse (cell-glass contact zone) for control and DZNep treated cells. Scale bars are 5 μm.

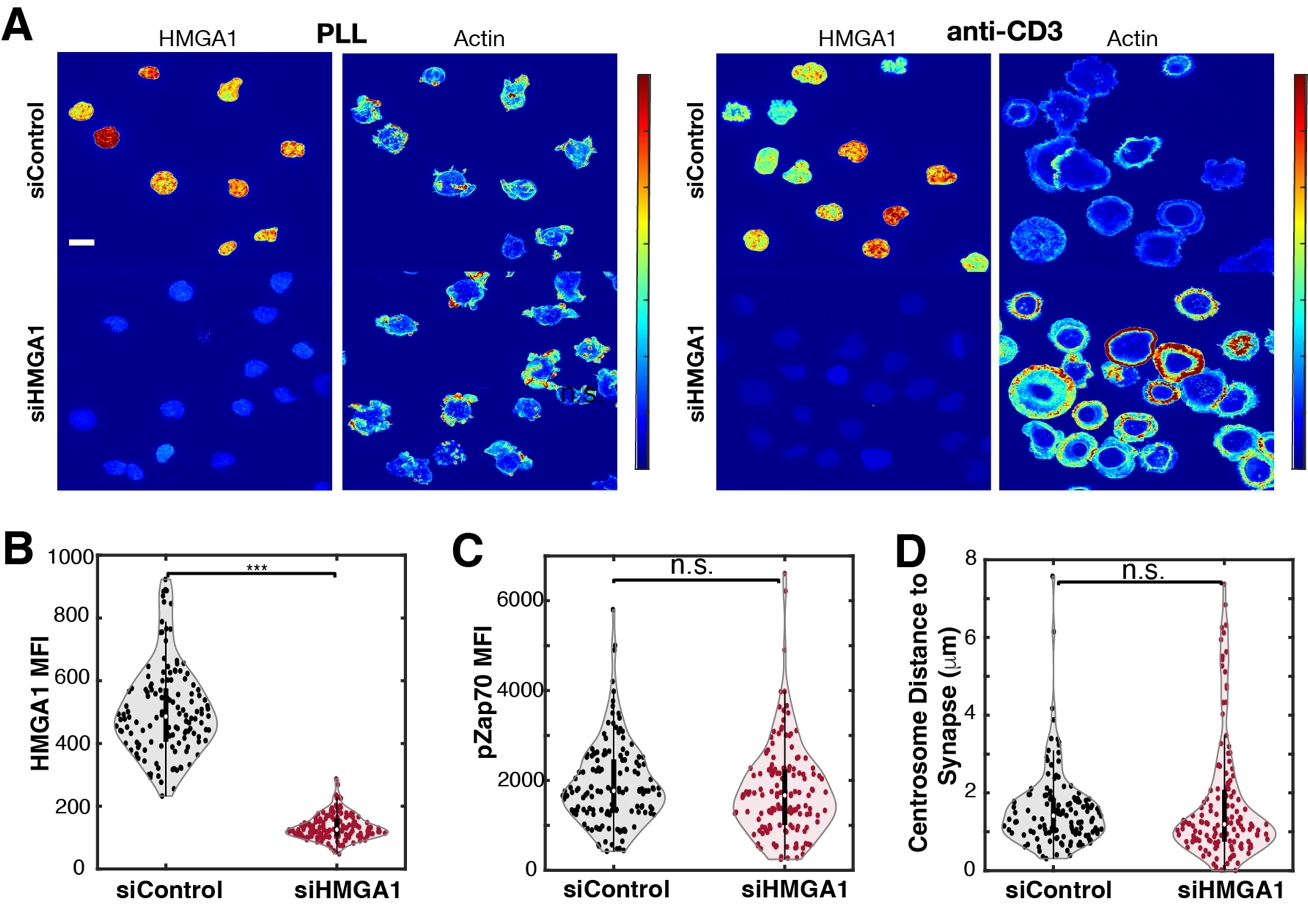

**Supplementary Figure 4. Chromatin decompaction by HMGA1 knockdown does not affect early TCR signaling.**

A) Maximum intensity *xy* projections of representative fields of view showing siControl and siHMGA1 cells plated on a PLL coated substrate (left) or on an anti-CD3 coated substrate (right). For each substrate, the images on the left panels correspond to immunostaining of HMGA1 and the images on the right panels to rhodamine-phalloidin staining of actin. B) Comparison of mean fluorescence intensity of HMGA1 immunofluorescence in siControl and siHMGA1 cells to validate HMGA1 knock down. C) Phosphorylated Zap70 mean fluorescence intensity in siControl and siHMGA1 cells. D) Centrosome distance to the synapse (cell-glass contact zone) in siControl and siHMGA1 cells. Scale bar is 10 μm.

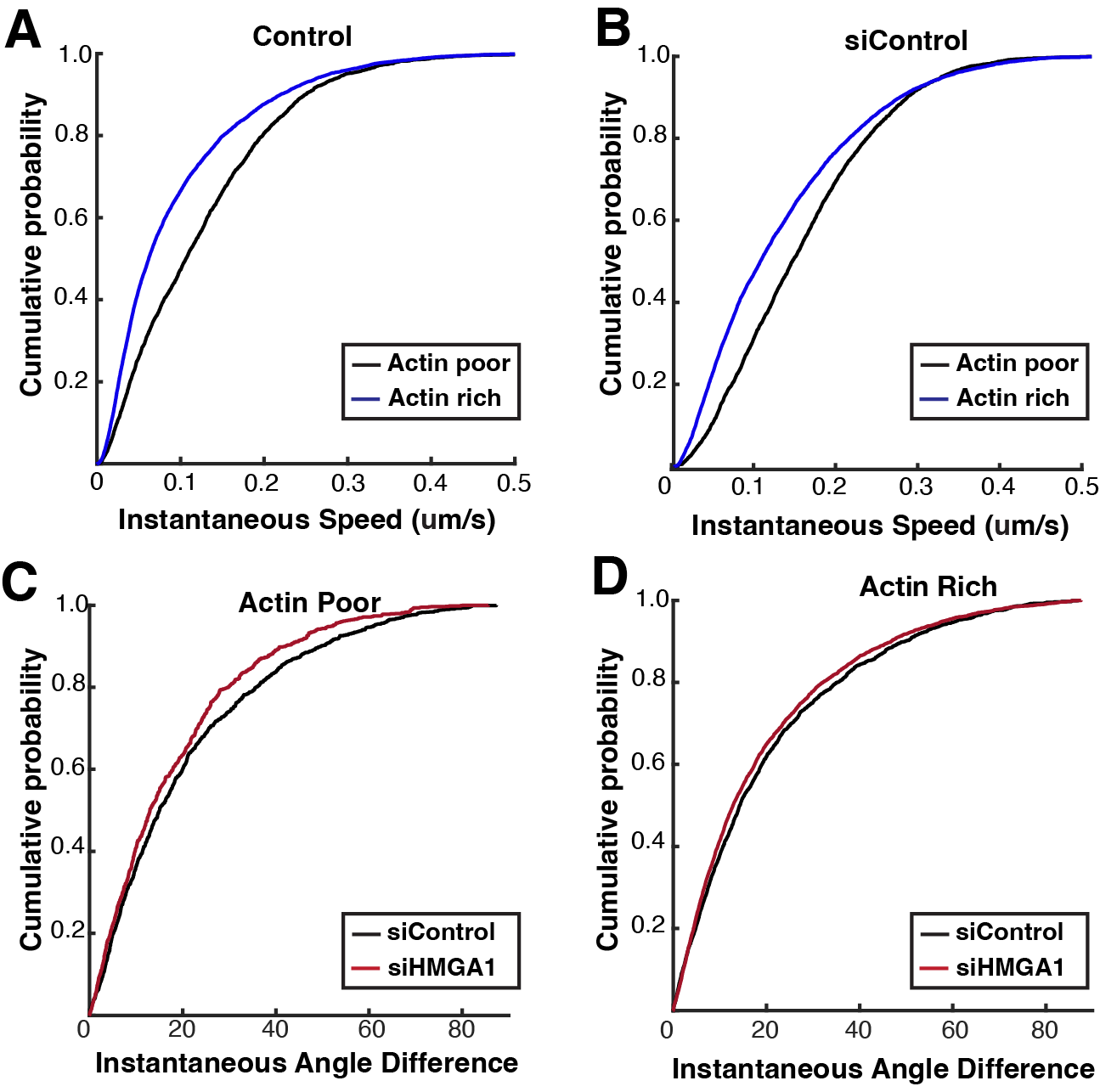

**Supplementary Figure 5. Effect of HMGA1 knockdown on EB3 speeds.**

Cumulative probability plots of EB3 instantaneous speeds in actin poor and actin rich regions for vehicle control (A) and siRNA control cells. (B). Cumulative probability plots of EB3 instantaneous angle difference in siControl and siHMGA1 cells at actin poor (C) and actin rich regions (D).

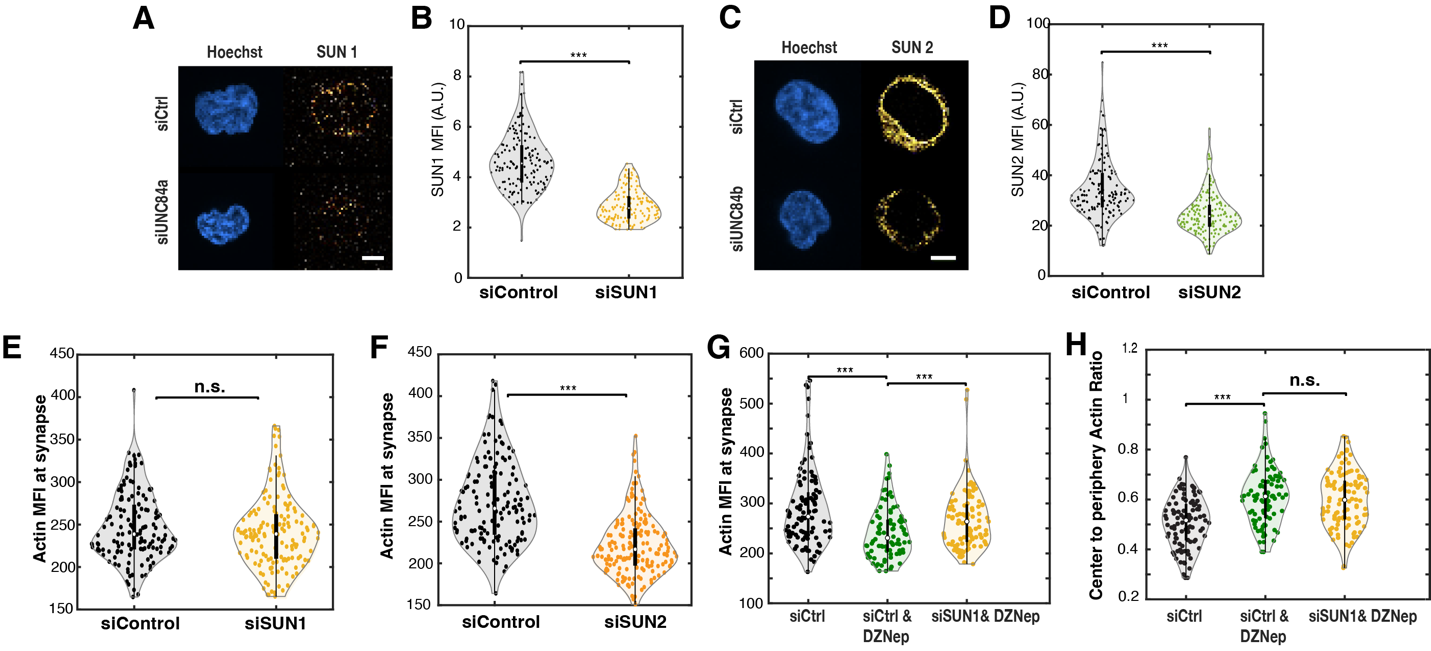

**Supplementary Figure 6. LINC complex proteins mediate bidirectional chromatin-actin cytoskeleton interactions.** A) Confocal images showing the distribution of SUN1 in siRNA control (siControl) and SUN1 knockdown (siSUN1) cells (right panels) and the nucleus labeled with Hoechst (left panels). B) Mean fluorescence intensity of SUN1 staining in siControl and siSUN1 cells. C) Confocal images showing the distribution of SUN2 in siRNA control (siControl) and SUN2 knock down (siSUN2) cells (right panels) and the nucleus labeled in Hoechst (left panels). D) Mean fluorescence intensity of SUN1 staining in siControl and siSUN2 cells. E) Mean fluorescence intensity of F-actin in siControl and siSUN1 cells. F) Mean fluorescence intensity of F-actin in siControl and siSUN2 cells. G) Mean fluorescence intensity of F-actin in siControl, siControl treated with DZNep and siSUN1 cells treated with DZNep. H) Ratio of central to peripheral actin for siRNA control (siControl), siControl treated with DZNep and siSUN1 cells treated with DZNep. Scale bars are 5 μm.

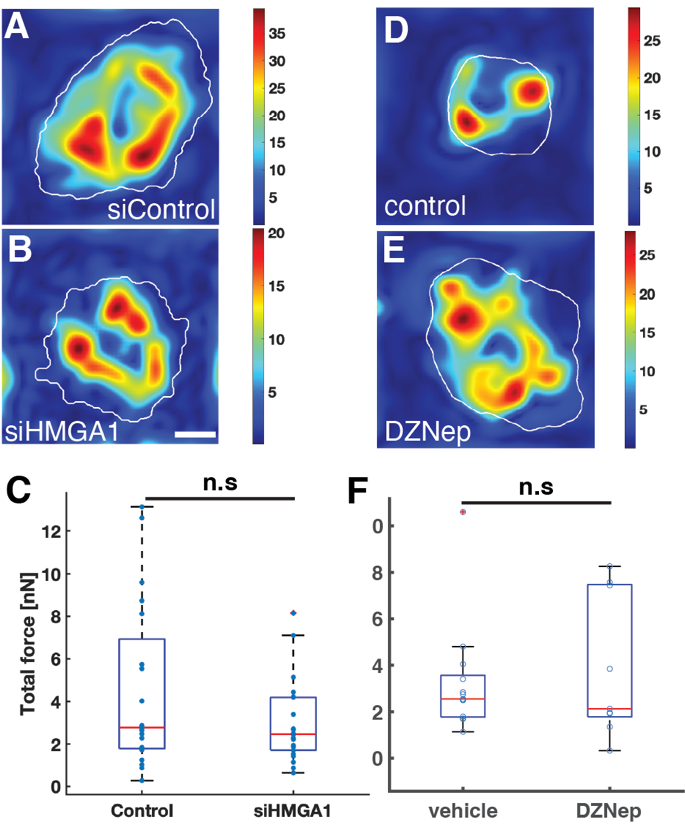

**Supplementary** **Figure** **7**. **Changes in chromatin compaction do not affect cellular force generation.** A-B) Representative traction force maps for siControl and siHMGA1 cells. Force scale is in Pa. C) Summary plot for total force exerted by cells (p=0.3; n=15 for siControl, 13 for siHMGA1). D-E) Representative traction force maps for vehicle and DZNep-treated cells. Force scale is in Pa. F) Summary plot for total force exerted by cells (p=0.4; n=12 for Control, 11 for DZNep). Scale bars are 5 μm.
